# Commitment in the PINK1/Parkin mitophagy decision circuit

**DOI:** 10.1101/2022.10.07.510659

**Authors:** Christopher S. Waters, Sigurd B. Angenent, Steven J. Altschuler, Lani F. Wu

## Abstract

Mechanisms that prevent accidental degradation of healthy mitochondria by the PINK1/Parkin mitophagy pathway are poorly understood. On the surface of damaged mitochondria, PINK1 accumulates and acts as the input signal to a positive feedback loop of Parkin recruitment, which in turn promotes mitochondrial degradation via mitophagy. However, PINK1 is also present on healthy mitochondria where it could errantly recruit Parkin and thereby activate this positive feedback loop. Here, we quantitatively mapped the relationship between PINK1 input levels and Parkin recruitment dynamics using live-cell microscopy and mathematical modeling. We found that Parkin is recruited to the mitochondria only if PINK1 levels exceed a threshold and only after a delay that is inversely proportional to PINK1 levels. The threshold and delay provide a “two-factor authentication” step for PINK1/Parkin activation. These properties arise from the PINK1/Parkin circuit topology and provide a mechanism for cells to assess damage signals before committing to mitophagy.

## Introduction

When mitochondria accumulate significant oxidative damage, they cease to function properly and can become depolarized. Cells maintain a healthy pool of mitochondria by degrading depolarized mitochondria via mitophagy (mitochondrial autophagy). A core molecular circuit mediating the mitophagy decision involves PINK1, a mitochondrial kinase, and Parkin, a cytoplasmic E3 Ubiquitin ligase (*1*). While PINK1 is constitutively recruited to the mitochondrial outer membrane (MOM), polarization-dependent degradation keeps PINK1 levels low on healthy mitochondria (*2*, *3*). Mitochondrial depolarization prevents this degradation and leads to PINK1 accumulation on the MOM (*2*, *3*). Upon PINK1 accumulation, positive feedback in the PINK1/Parkin circuit leads to extensive labeling of the mitochondrial surface with phospho-Ubiquitin (pUb) and to pUb-dependent recruitment of downstream mitophagy machinery (*1*, *4*, *5*).

Positive feedback in the PINK1/Parkin circuit proceeds through several steps: PINK1 phosphorylates mitochondrial Ubiquitin, pUb recruits autoinhibited Parkin to the MOM, PINK1 phosphorylates Parkin’s Ubiquitin-like (UBL) domain to activate Parkin, and activated Parkin deposits new Ubiquitin on proteins at the MOM (*1*, *6*–*14*). Newly deposited Ubiquitin can then be phosphorylated by PINK1, starting new rounds of amplification (*1*, *15*). Together, these steps form a PINK1-dependent positive feedback circuit topology, where PINK1 provides an input signal that regulates continued recruitment of Ubiquitin and Parkin (Fig. 1). Prior work speculates that the PINK1/Parkin feedback topology plays a role in preventing errant mitophagy decisions (*4*, *6*, *15*–*17*). Activation thresholds and other emergent properties have been speculated to confer tolerance to noise, such as transient increases in mitochondrial PINK1 (*6*, *15*–*18*). However, in practice, it remains unclear how noise tolerance arises within the PINK1/Parkin positive feedback circuit.

**Fig. 1.**
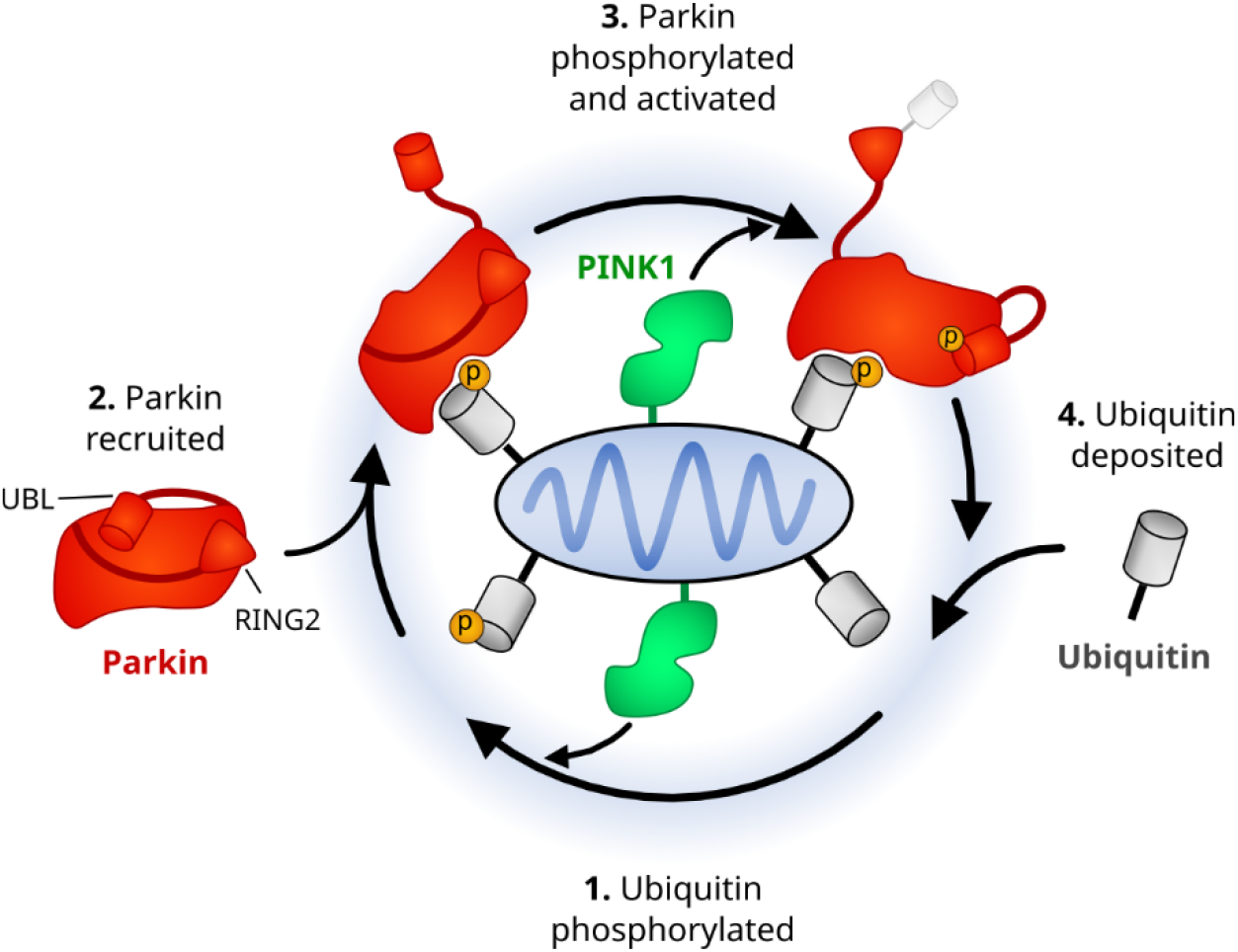
PINK1 provides continual input to the PINK1/Parkin positive feedback loop. Cartoon schematic of the PINK1/Parkin positive feedback loop. Phosphorylation sites, yellow circles. Parkin’s Ubiquitin-like (UBL) domain and catalytic RING2 domain labeled to illustrate conformational changes required for Parkin’s activation. In each round of amplification, 1) PINK1 phosphorylates Ubiquitin, 2) pUb recruits autoinhibited parkin from the cytoplasm to the mitochondria, freeing Parkin’s UBL domain, 3) PINK1 phosphorylates Parkin’s UBL domain, freeing the catalytic RING2 domain and activating Parkin, 4) activated Parkin deposits additional Ubiquitin. Ubiquitin is attached to proteins on the MOM to form Ubiquitin chains. Ubiquitin chains not shown.

Here, we investigated how cells avoid accidental degradation of healthy mitochondria by PINK1 and Parkin. We used live-cell microscopy to quantitatively map the relationship between discrete PINK1 levels and Parkin recruitment dynamics, revealing inherent regulatory properties that allow the PINK1/Parkin mitophagy pathway to only activate in the presence of both elevated and sustained mitochondrial PINK1 levels. We found that both of these properties can be modulated by inhibiting the mitochondrial deubiquitinase USP30, an antagonist of the PINK1/Parkin circuit (*4*, *19*–*21*). Mathematical modeling then revealed that the topology of the PINK1/Parkin circuit, alone, is sufficient to generate the observed regulatory properties.

## Results

To study the PINK1/Parkin circuit, we examined the relationship between circuit input (mitochondrial PINK1 levels) and activation (mitochondrial Parkin recruitment) on healthy mitochondria. We designed an experimental system that would: avoid confounding side effects from chemically induced mitochondrial depolarization (e.g., dysregulation of ATP, reactive oxygen species (ROS), and calcium homeostasis) (*22*–*26*), enable precise control of mitochondrial PINK1 input levels, and allow simultaneous monitoring of circuit input and activation at a single-cell level over time.

### Building a fluorescent Parkin probe to read out circuit activation

First, we focused on establishing methods to monitor circuit activation. A long-established readout of PINK1/Parkin circuit activation and of mitophagy initiation is Parkin relocalization from the cytoplasm to the mitochondria (*2*). This relocalization is commonly monitored through N-terminally tagged Parkin fusions. However, N-terminal epitope tags have been reported to disrupt Parkin’s autoinhibited state, presumably by weakening interactions involving Parkin’s N-terminal UBL domain (*27*), a key domain for both Parkin’s autoinhibition and activation (*10*, *11*, *13*) (Fig. 1; Fig. 2A).

**Fig. 2.**
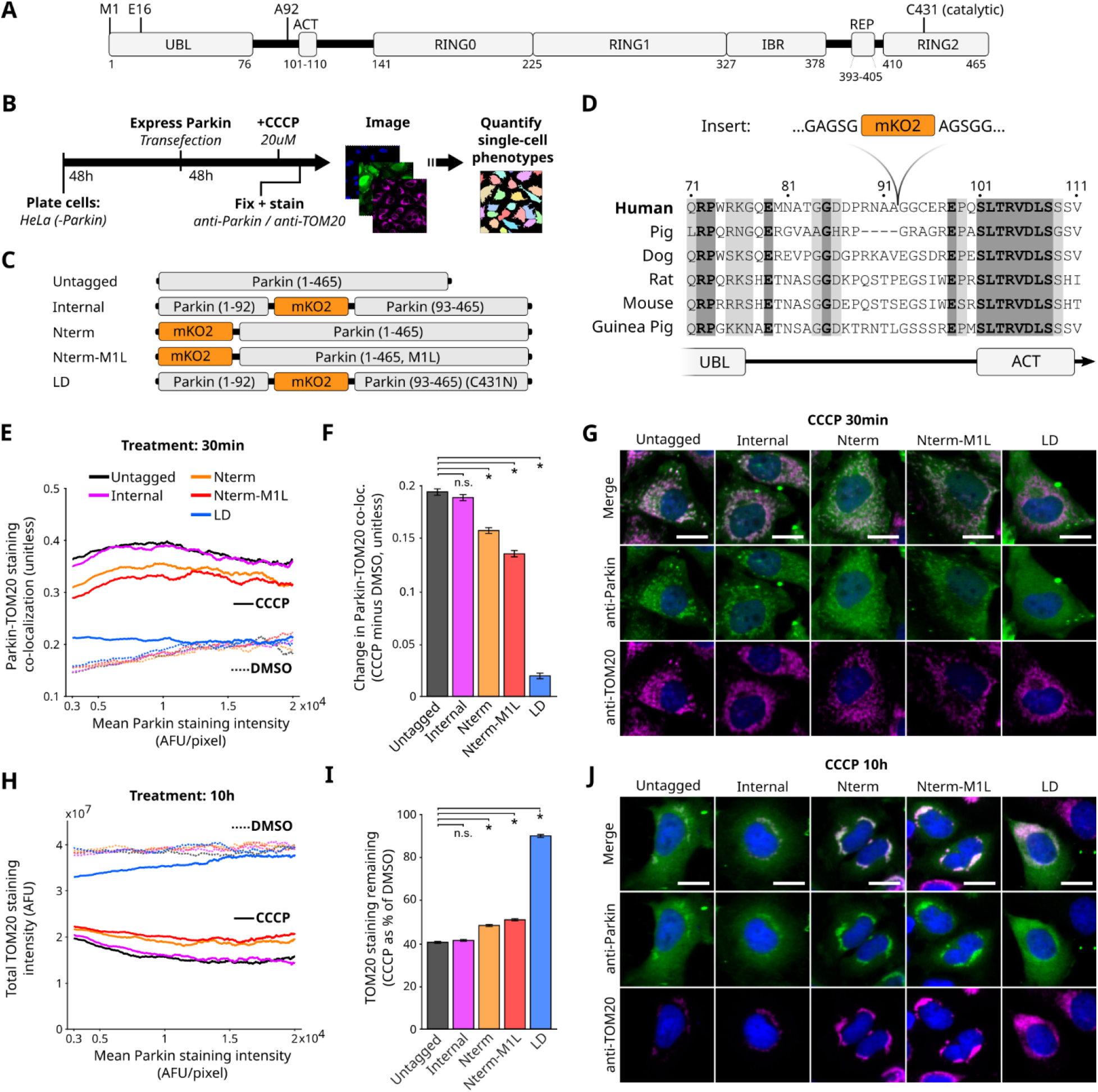
Identification of an internal Parkin tagging site that preserves function. **(A)** Domains and selected residues of Parkin. **(B)** Immunofluorescence (IF) approach for assessing Parkin function. HeLa cells lack endogenous Parkin expression. **(C)** Parkin fusion proteins, assessed for activity. LD, ligase dead. **(D)** Conservation of residues surrounding Parkin’s internal tagging site. **(E)** Parkin recruitment to mitochondria (TOM20) as a function of Parkin expression. Data are sliding medians of single-cell data shown in Fig. S2A. AFU, arbitrary fluorescence units. **(F)** Mean change in Parkin localization across Parkin expression levels from (E). Data are mean and 95% confidence intervals, calculated via bootstrap analysis. *adjusted p-value<0.05 by bootstrap analysis and Bonferroni multiple comparison adjustment. **(G)** Representative images showing Parkin recruitment. Scale bars, 20um. **(H)** Total mitochondrial intensity as a function of Parkin expression. Data are sliding medians of single-cell data shown in Fig. S2B. **(I)** Mean mitochondrial depletion across Parkin expression levels. Representation and analysis as in (F). **(J)** Representative images showing mitochondrial loss. Scale bars, 20um.

What might cause N-terminal tag-based Parkin dysfunction? The N-terminus of Parkin’s UBL domain has not been reported to be involved in intra-or inter-protein interactions (*12*–*14*, *28*). However, we observed that the positively charged amino group at Parkin’s N-terminus is an integral part of the UBL domain fold: the NH_3_^+^ group of Methionine 1 (M1) can form a salt bridge with the negatively charged sidechain of glutamic acid 16 (E16) on an adjacent beta strand (Fig. S1A-C; methods). We observed similar salt bridges in structures for other N-terminal UBL domains and for Ubiquitin itself (Fig. S1D-F). Furthermore, alignment of all known UBL domain sequences revealed that negatively charged residues are conserved at an equivalent position to E16 in N-terminal, but not internal, UBL domains (Fig. S1G). Finally, we found that a Parkin (E16A) mutant displayed both reduced mitochondrial recruitment and reduced downstream mitochondrial degradation following mitochondrial depolarization (Fig. 2B; Fig. S2H). Together, these data suggest that the conserved NH_3_^+^-E16 interaction in Parkin’s UBL domain is functionally important and that N-terminal tags may disrupt this interaction by replacing the NH3^+^ group.

To engineer a fluorescently tagged Parkin with preserved activity, we investigated alternative tagging locations. C-terminal tags were not pursued both due to proximity to Parkin’s catalytic RING2 domain and due to involvement of Parkin’s C-terminus in intra-protein interactions (*14*) (Fig. S1A). Instead, we identified an internal tag insertion site, between A92 and G93, within a disordered (*12*, *13*) and non-conserved linker region (Fig. 2A,C-D). We then tested a panel of tagged Parkin fusions with monomeric Kusabira-Orange 2 (mKO2; Fig. 2B-C). As expected, N-terminally tagged Parkin displayed reduced recruitment to and degradation of mitochondria (Fig. 2E-J; Fig. S2). An M1L substitution, present in commonly used N-terminal Parkin fusions (*29*), further decreased mitophagy activity. Internally tagged Parkin, on the other hand, displayed mitophagy activity similar to untagged Parkin. We confirmed that recruitment of internally tagged Parkin to depolarized mitochondria was reversable upon mitochondrial repolarization, a well-established property of the circuit (*2*, *3*, *30*). Based on our results, we chose to monitor activation of the PINK1/Parkin circuit using internally tagged Parkin (hereafter, Parkin-mKO2).

### Control of circuit input with simultaneous monitoring of circuit activation

Next, we established methods to activate and monitor circuit input by developing the ability to directly control the amount of PINK1 on mitochondria in HeLa cells stably expressing Parkin-mKO2. We adapted existing approaches, allowing chemically induced recruitment of PINK1 to mitochondria (*2*, *31*–*33*), for quantitative live-cell microscopy (Fig. 3A-C). Specifically, the FRB-FKBP rapamycin analog (rapalog) induced heterodimerization system was used (*31*, *33*). PINK1’s mitochondrial targeting transmembrane region (MTS; amino acids 1-110) was replaced with an FKBP domain and a fluorescent mNeonGreen tag was added (PINK1-mNG; Fig. 3A-C). Additionally, an FRB domain tether was fused to a MOM-targeting domain and to a SNAP tag for far-red live-cell imaging via staining with SNAP-Cell 647-SiR (MtTether-SNAP + SNAP-647; Fig. 3A-C). PINK1-mNG was expressed using a doxycycline-inducible promoter, while MtTether-SNAP was stably expressed and localized to the mitochondria (Fig. S3A-C). We verified that addition of 200nM rapalog causes the expected sequence of: PINK1-mNG recruitment to mitochondria, Parkin-mKO2 recruitment to mitochondria, and mitochondrial aggregation/degradation (Fig. 3D; Fig. S3C-D). We further observed that: Parkin-mKO2 was not recruited in the absence of rapalog (Fig. S3B/D; Fig. 3D), mitochondria did not depolarize in response to PINK1-mNG or Parkin-mKO2 recruitment (data not shown), and mTOR signaling was not affected by rapalog treatment (Fig. S3E). We selected 12 hours post rapalog addition as our final imaging time point to give cells sufficient time to undergo Parkin-mKO2 recruitment. Finally, cell-to-cell heterogeneity of PINK1-mNG expression (Fig. S3A-B) allowed us to simultaneously observe a range of mitochondrial PINK1-mNG levels in each imaging experiment. Thus, our system allowed us to control mitochondrial PINK1-mNG input and to simultaneously monitor mitochondrial PINK1-mNG and Parkin-mKO2 levels via live-cell microscopy.

**Fig. 3.**
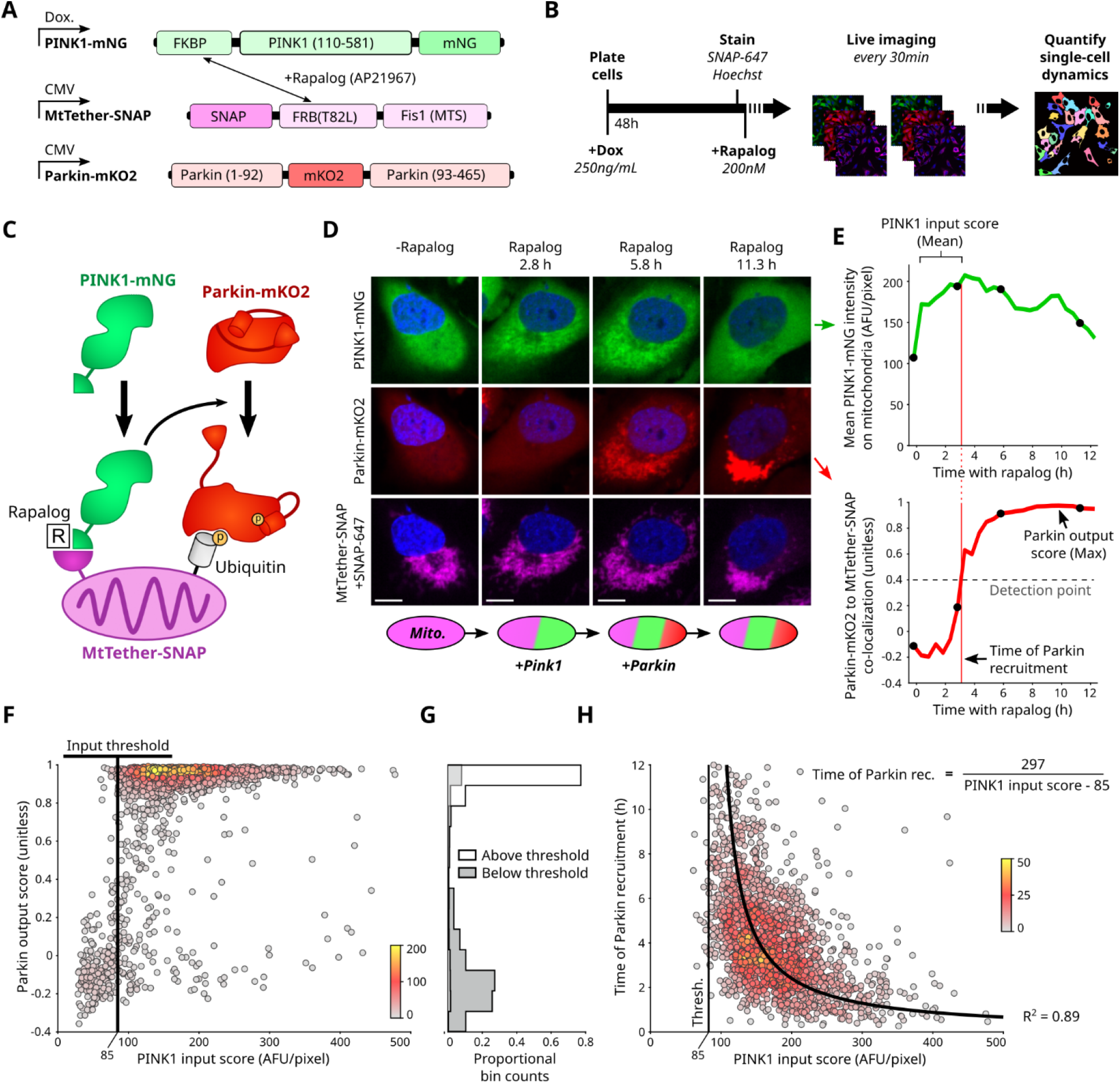
Input requirements for activation of the PINK1/Parkin circuit. **(A)** Rapalog-induced PINK1 recruitment system. MTS, mitochondrial targeting sequence. T82L, mutation required for rapalog binding. Expression methods: Doxycycline-inducible (dox.) or cytomegalovirus (CMV) constitutive promoters. **(B)** Live-cell imaging approach. **(C)** Cartoon of induced recruitment system **(D)** Representative images showing PINK1-mNG and Parkin-mKO2 recruitment to mitochondria. Scale bars, 10um. **(E)** Quantification of PINK1/Parkin dynamics for the cell shown in (D). Co-localization, intensity correlation. Black dots, timepoints shown in (D). **(F)** Parkin output scores versus PINK1 input scores for 1987 individual cells. Colors: local point density (number of nearby cells, methods). **(G)** PINK1 input threshold separates Parkin activated from Parkin non-activated cells. **(H)** Parkin recruitment time versus PINK1 input scores for 1676 individual cells. Colors: local point density (methods). Fitted hyperbolic curve and R^2^ value are shown.

### Quantification of circuit dynamics

Using the rapalog-induced recruitment system, we defined an approach to quantify circuit dynamics at single-cell resolution by tracking mitochondrial PINK1-mNG levels (*via* mean PINK1-mNG intensity) and Parkin-mKO2 recruitment (*via* Pearson correlation of Parkin-mKO2 and MtTether-SNAP intensities) (Fig. 3D-E; methods). Each cell was assigned: a PINK1 input score (time-averaged mitochondrial PINK1-mNG levels before Parkin-mKO2 recruitment; methods, Fig. 3E, top); a time of circuit activation defined by when Parkin recruitment is first observed (Parkin-MtTether intensity correlation > 0.4; methods; Fig. 3E, bottom); and a Parkin output score (maximum observed level of Parkin-mKO2 recruitment; Fig. 3E, bottom). This triplet of scores summarized circuit behavior for each cell.

### PINK1 input threshold determines decision for Parkin recruitment

The input-to-output relationship in this PINK1-Parkin circuit was bimodal, having either undetectable or high Parkin responses (Fig. 3F; bimodality was also observed using other scores; Fig. S4). Furthermore, a threshold for PINK1 input scores effectively separated these low *vs* high Parkin output cells (Fig. 3F-G; methods). These data suggest that the PINK1-Parkin circuit displays switch-like behavior, where circuit activation is triggered if input levels surpass an input threshold.

### PINK1 levels determine timing of Parkin recruitment

Next, we examined how circuit input scores influence the detection time of Parkin recruitment. We found that the relationship between single-cell PINK1 input scores and time to detection of Parkin response was well fit by a hyperbolic model (R-squared value of 0.89; Fig. 3H; a linear model is inconsistent with the observed threshold behavior). That is, we observe an “input reciprocal delay” behavior: doubling the amount by which PINK1 input exceeds the threshold will halve the time to trigger a Parkin response. Both the input threshold and input reciprocal delay required of PINK1 levels for Parkin recruitment were recapitulated by fixed-cell time series (Fig. S5). Together, the input threshold and input reciprocal delay behaviors describe a “two-factor” regulatory paradigm for the PINK1/Parkin pathway.

### Two-factor authentication can be perturbed

We next tested how chemical perturbations reported to sensitize or desensitize the PINK1/Parkin pathway, in depolarization-based assays, would affect the two activation requirements identified above (Fig. 4A-B). To reduce pathway sensitivity, we tested the ROS scavenger N-acetyl-L-cysteine (NAC) (*22*). ROS generated by depolarized mitochondria is thought to promote mitophagy (*23*). We found that treatment with 2mM NAC delayed Parkin recruitment but did not alter the input threshold (Fig. 4C; Fig. S6). The direction of this effect is consistent with previously described effects of NAC treatment (*22*) and confirms that NAC treatment can directly decrease PINK1/Parkin circuit sensitivity, even in the absence of overt mitochondrial depolarization, via an unknown mechanism.

**Fig. 4.**
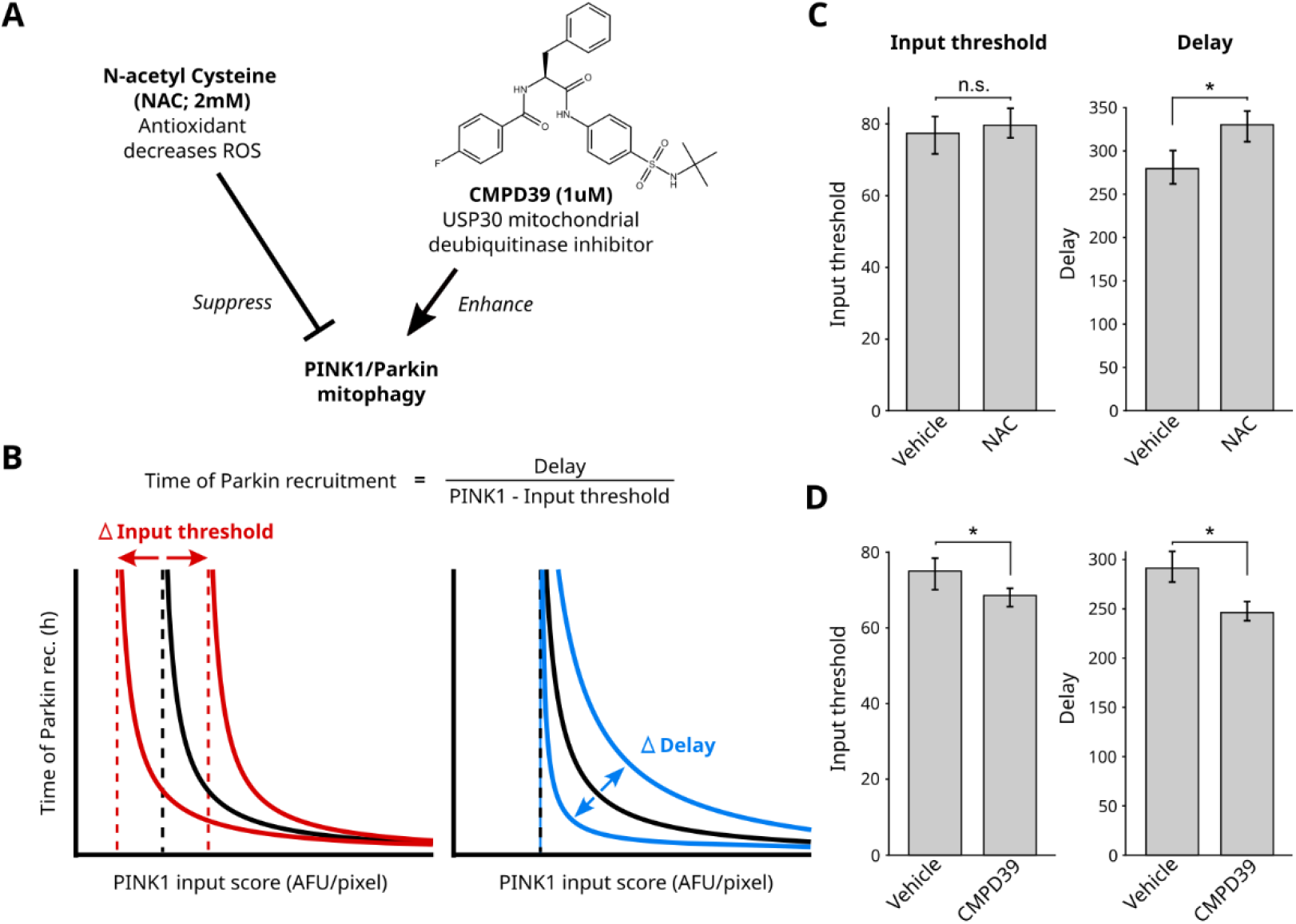
Dissecting the effects of mitophagy modifiers on PINK1/Parkin activation. **(A)** Reported effects of mitophagy perturbations from the literature. Chemical structure of the USP30 inhibitor, CMPD39. **(B)** Potential effects of perturbations on circuit behavior. **(C)** Measured threshold and delay values for live cells in NAC or vehicle conditions. Data are mean and 95% confidence intervals calculated via bootstrap analysis. *adjusted p-value<0.05 by bootstrap analysis and Bonferroni multiple comparison adjustment. Single-cell data shown in Fig. S6. **(D)** Quantified values for USP30 inhibition as in (C). Single-cell data shown in Fig. S7.

To increase pathway sensitivity, we tested the USP30 inhibitor CMPD39 (*34*). USP30 is thought to both limit the basal mitochondrial Ubiquitin level (*6*, *16*, *17*, *35*) and directly antagonize PINK1/Parkin circuit activation (*4*, *19–21*) through removal of mitochondrial Ubiquitin. In agreement with these roles, we found that treatment with CMPD39 both shortened the Parkin recruitment delay and lowered the input threshold (Fig. 4D; Fig. S7).

### Input-coupled positive feedback is sufficient to produce two-factor authentication

We next investigated emergent behaviors of the PINK1/Parkin circuit topology using mathematical models (simplified model: Fig. 5; generalized model and analysis: Fig. S8, Fig. S9, supplementary text). A defining feature of these models is that the input controls the feedback strength, which we refer to as “input-coupled positive feedback” (Fig. 5A). When PINK1 levels are zero, the circuit is unable to become activated and is in a stable “off” state, as expected because both Parkin activation and Parkin recruitment depend on PINK1 phosphorylation activity. If PINK1 levels increase above an input threshold, positive feedback for Parkin recruitment will be initiated due to a transcritical bifurcation arising within the system (Fig. 5B; Fig. S8A-C; supplementary text) (*36*–*39*). Finally, if an on-state level of Parkin is defined (e.g., high enough for detection or for recruitment of downstream mitophagy components), then mathematical analysis shows that the time to reach this on-state is inversely related to PINK1 levels (Fig. 5C; Fig. S8D-E; supplementary text). Simulations using cell-to-cell heterogeneity in model parameters (methods) yielded a distribution of responses along the analytical solutions and qualitatively matched experimental data (Fig. 5B-C; Fig. S4A; Fig. 3H). Thus, our models recapitulated the experimentally observed noise-tolerance properties of the PINK1/Parkin circuit.

**Fig. 5.**
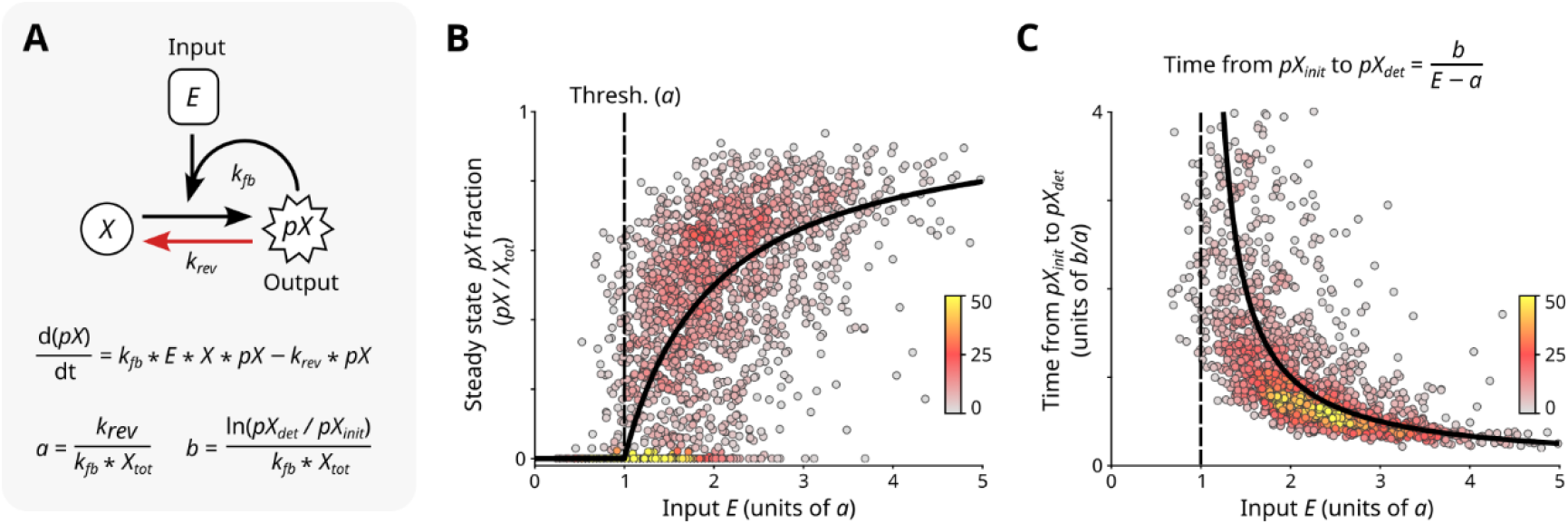
Threshold and delay properties arise within a minimal model of input-coupled positive feedback. **(A)** Minimal model of input-coupled positive feedback. Total concentration of *X: X_tot_* = *X* + *pX*; initial concentration of *pX*: *pX_init_* > 0; detection concentration of *pX*: *pX_det_* << *X_tot_*. Parameters *a* and *b* govern input threshold and reciprocal time delay, respectively. Derivations and model generalization described in supplemental text and Fig. S8. **(B-C)** Steady state analysis (B) and relationship between input and time to reach detectable output levels (C). Black curves: algebraic solutions of (A). Points: simulated responses for heterogeneous single-cell values of *E, k_rev_*, and *pX_init_* (n = 2000 points; methods). Colors: local point density (methods). Axes correspond to data: for (B) in Fig. S4A or (C) in Fig 3H.

## Discussion

This study reveals an intrinsic mechanism by which the PINK1/Parkin circuit can avoid errant activation. We developed an approach to modulate and measure mitochondrial PINK1 input levels while simultaneously monitoring Parkin localization responses using a novel tagging strategy. Our input-to-response mapping confirms previous speculation that a PINK1 threshold buffers against improper activation of the PINK1/Parkin pathway. However, our analysis also reveals that Parkin activation does not simply follow from the requirement that PINK1 levels exceed a threshold. Rather, appreciable Parkin activation is delayed for a duration of time that is inversely proportional to PINK1 levels. The dependence of circuit activation on persistence of input signal could allow mitochondria to ignore transient depolarizations yet respond to persistent depolarization or to repeated transient depolarizations (*30*). These regulatory properties can be modulated to tune the sensitivity of the pathway. Finally, mathematical modeling shows that input-coupled positive feedback—central to the PINK1/Parkin circuit topology—can intrinsically give rise to input thresholds and input reciprocal delay behaviors that enable noise-tolerance.

This study opens several questions for future investigation. First, while we confirm that an activation threshold for the PINK1/Parkin circuit exists and can be controlled by USP30 (*6*, *15*–*17*), our data does not resolve whether USP30 controls this threshold by removing Parkin-deposited Ubiquitin or basal mitochondrial Ubiquitin. Second, a number of additional proteins have been implicated in controlling mitophagy. These proteins include the putative Ubiquitin phosphatase PTEN-L (*40*), deubiquitinases such as USP15 (*41*), and Parkin-binding proteins such as Miro (*25*). It remains unclear how each of these proteins modulates PINK1/Parkin dynamics within the conceptual framework described here. Finally, depolarization-dependent signals including ROS generation, calcium release, and ATP depletion (*22*–*26*) have been associated with modulating PINK1/Parkin activation sensitivity. However, the mechanisms by which these signals affect the PINK1/Parkin pathway have been difficult to disentangle using depolarization-based assays (*22*). The quantitative framework described in this work provides a starting point for understanding how various proteins and signals affect properties of PINK1/Parkin circuit dynamics.

Input-coupled feedback circuits provide control over both when and where amplification can occur. For example, because the presence of input is required to maintain activation, PINK1/Parkin circuit activation is not expected to spread to healthy mitochondria where PINK1 is not present at activating levels. When positive feedback is decoupled from input, as in many classic feedback systems, spatially constrained activation may be lost without additional mechanisms (*36*, *38*). Beyond mitophagy, there are other systems where input-coupled positive feedback may operate. In EGF receptor (EGFR) signaling, an EGFR-driven positive feedback loop involving GAB1, PI3K and PIP3 matches the topology of an input coupled positive feedback circuit (*42*, *43*). A similar positive feedback loop also exists for insulin receptor (IR) signaling via IRS, PI3K, and PIP3 (*43*). Our work motivates future studies into how input-coupled positive feedback circuits enable different cellular-decision making strategies in space and time.

---

**Fig. S1.**
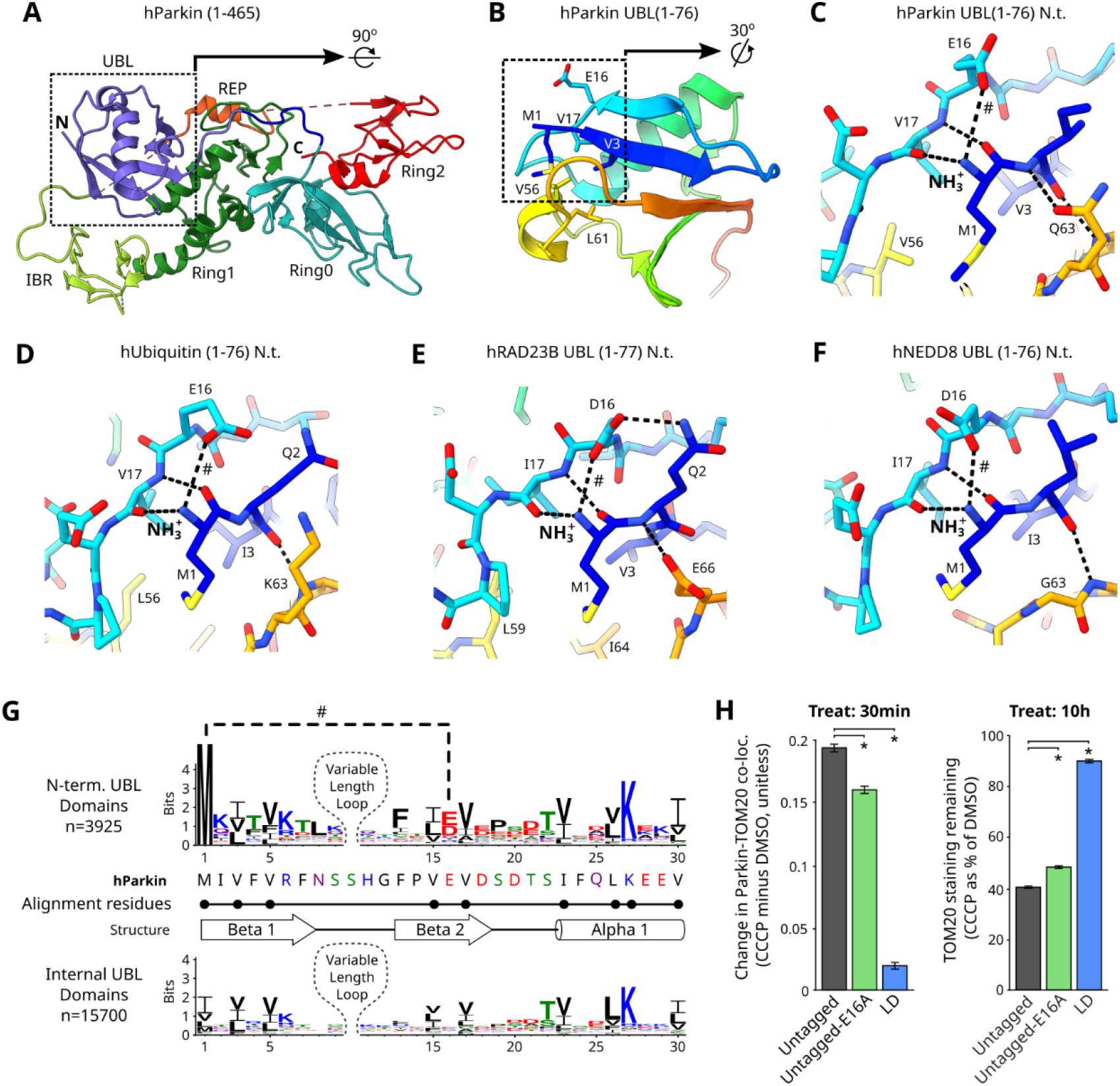
Structural importance of Parkin’s N-terminus. **(A-C)** Human Parkin structure. **(A)** Full protein, colored by domain. **(B)** UBL domain colored blue to red, N to C terminus. **(C)** N-terminus (N.t.) of UBL domain. Carbons colored as in B; Oxygen, red; Nitrogen, blue; sulfur, yellow. Dotted lines, predicted interactions. # indicates predicted salt bridge between E16 and the N-terminal NH_3_^+^, in all panels. **(D-F)** N.t. structures of Ubiquitin and other N-terminal UBL domains. **(G)** Sequence logo showing amino acid conservation in UBL domains. Analysis across unique UBL domains in the Uniprot database. Residues used for motif-based alignment are marked. **(H)** Functional quantification of untagged Parkin with an E16A mutation as in Fig. 2E-J. Data collected and analyzed alongside data in Fig. 2E-J. Data are mean and 95% confidence intervals calculated via bootstrap analysis. *adjusted p-value<0.05 by bootstrap analysis and Bonferroni multiple comparison adjustment. LD, ligase dead.

**Fig. S2.**
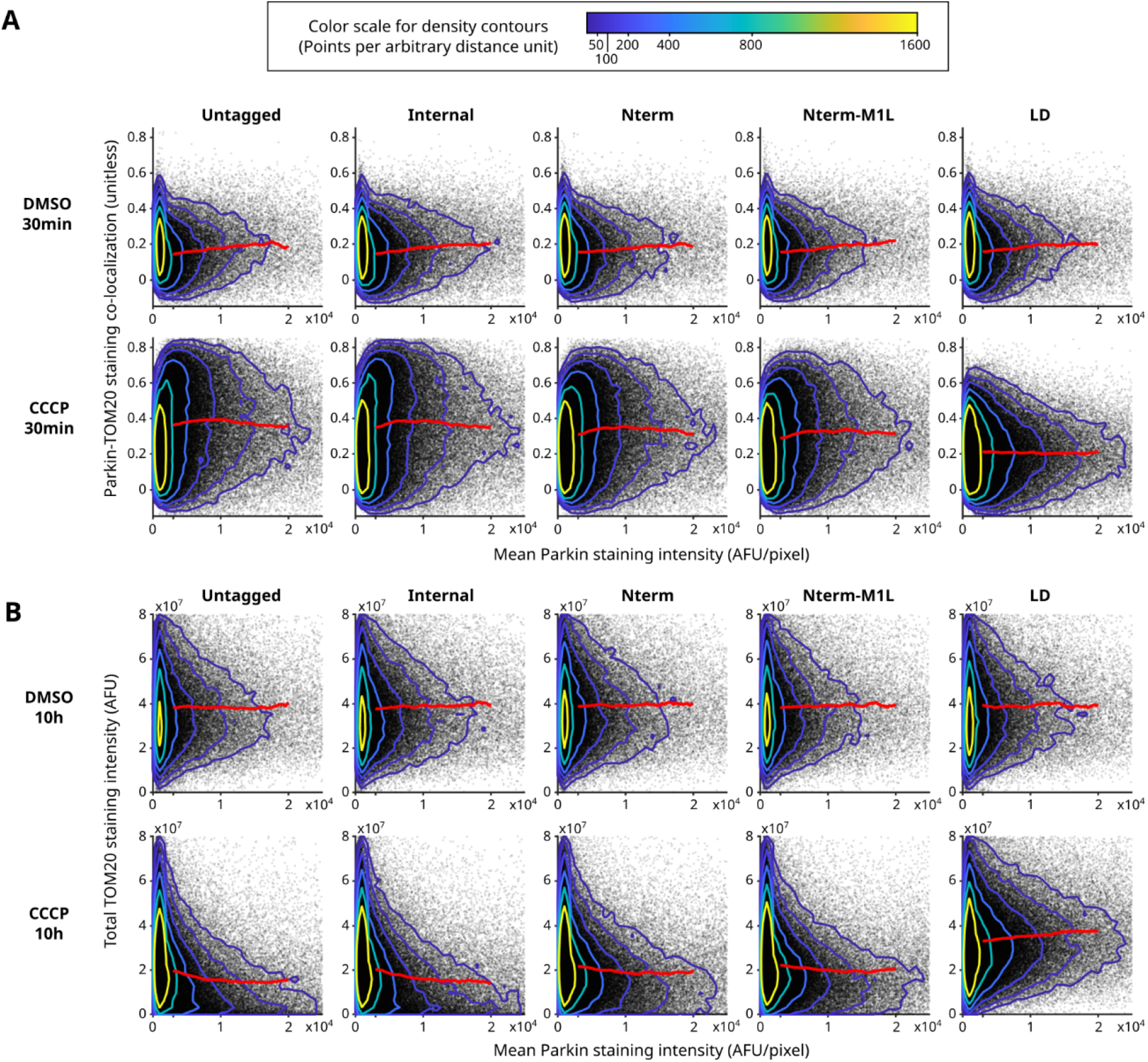
Comparison of Parkin tagging strategies. **(A & B)** Single-cell data used to generate Fig. 2E-J. Points, single cells; contours, datapoint density; red lines, sliding medians. Single-cell data from three experimental replicates was pooled. DMSO: n=~60,000 cells/condition; CCCP: n=~150,000 cells/condition; ~35% of cells had Parkin expression within the range used to calculate the moving medians, 3000-20000 AFU/pixel.

**Fig. S3.**
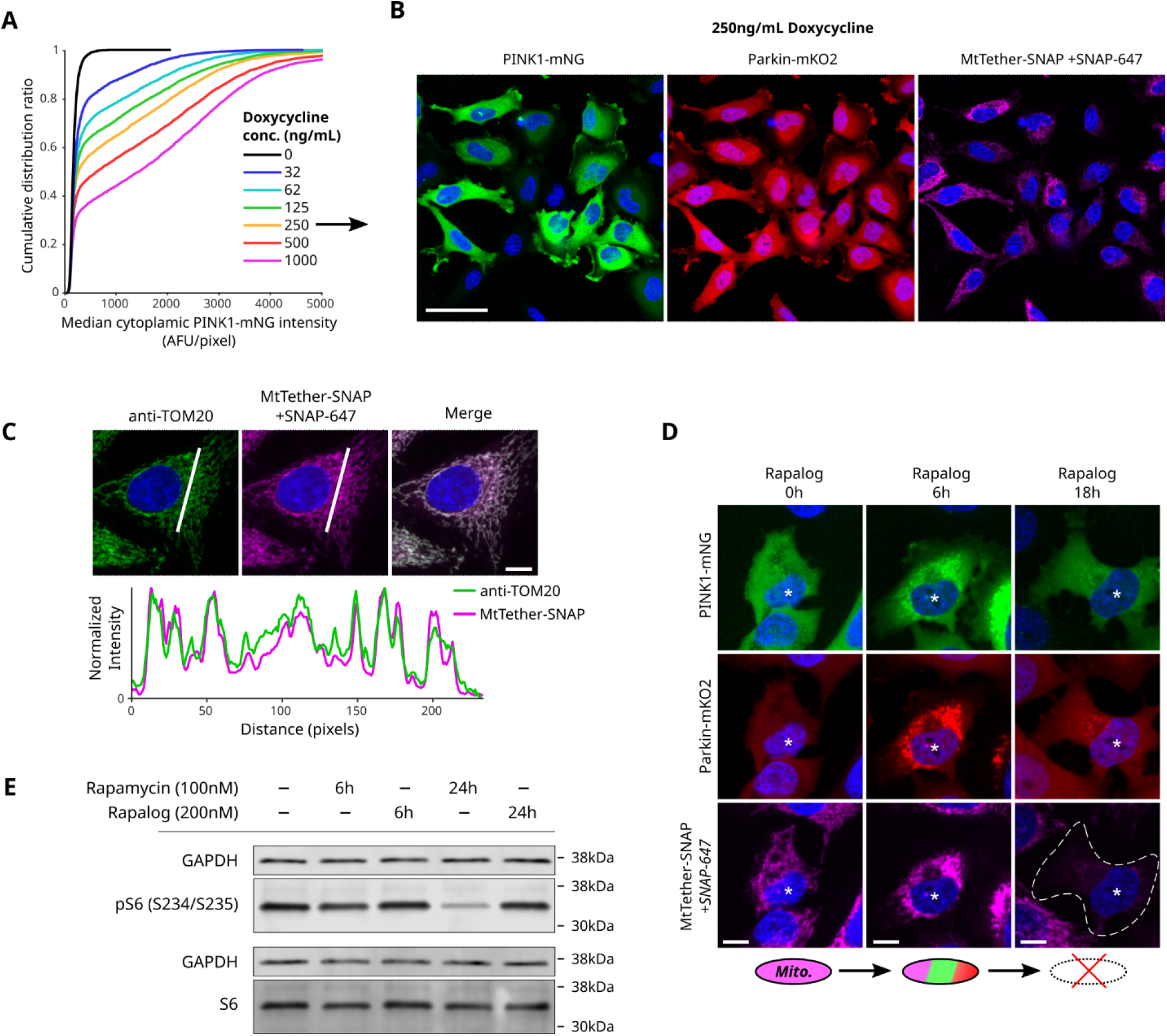
Characterization of the inducible PINK1/Parkin circuit. **(A)** Cumulative distribution of single-cell PINK1 expression levels following 48h doxycycline treatment. **(B)** Representative images showing population-level expression levels. Scale bar, 50um. **(C)** Top: Representative cell showing colocalization between MtTether-SNAP stained with SNAP-647 and mitochondria immunostained for TOM20. Bottom: pixel intensity values along white line in the single-channel images above. Scale bar in merged channel image, 20um. **(D)** Representative images showing mitochondrial loss following prolonged rapalog-driven PINK1-mNG and Parkin-mKO2 recruitment. Asterisk indicates cell of interest. Dotted line in bottom right image indicates cell boundary. Scale bars, 10um. **(E)** Western blot comparison of phosphorylated S6 (top pair) to unphosphorylated S6 (bottom pair) following treatment with either rapamycin or rapalog.

**Fig. S4.**
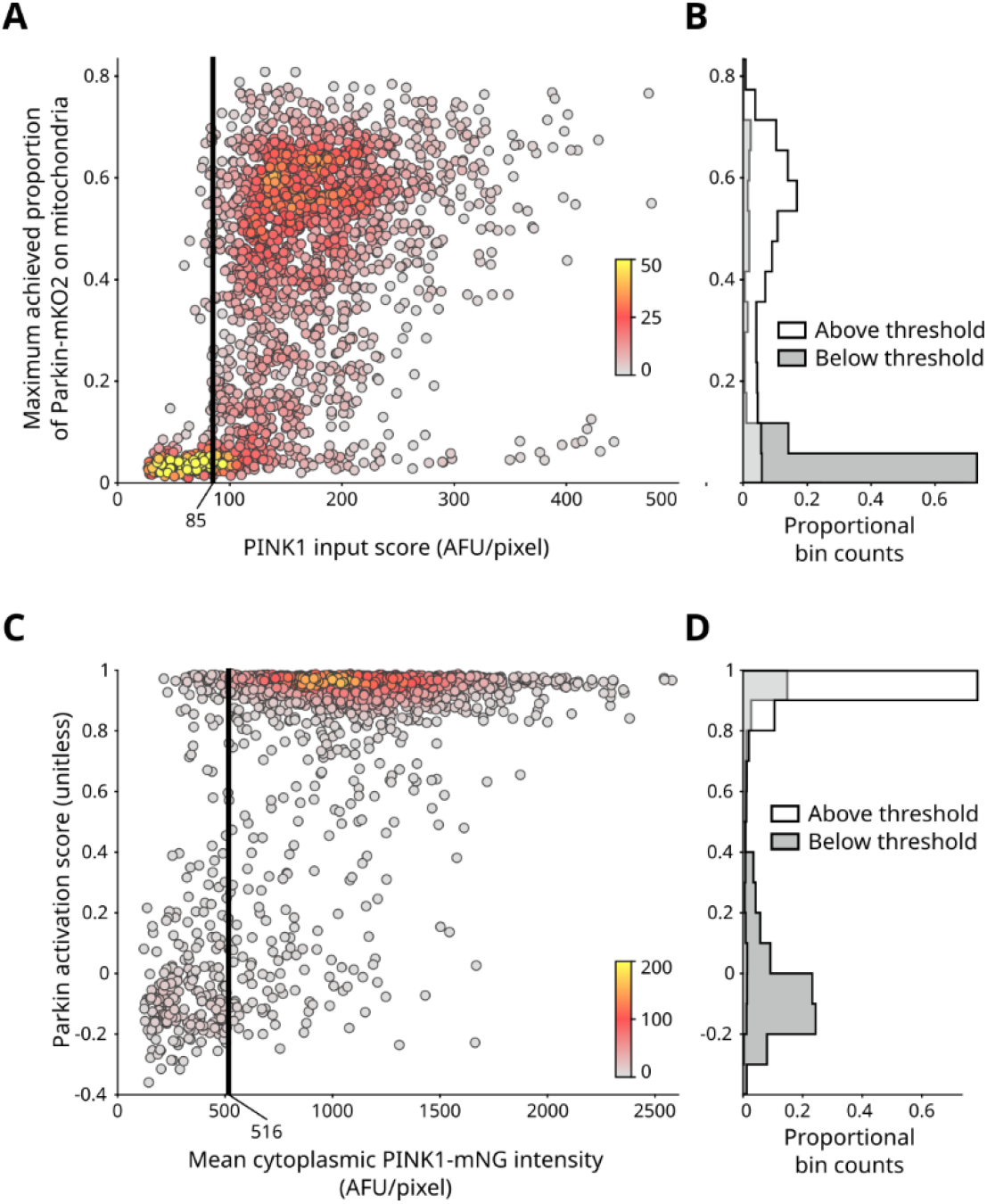
Alternative PINK1/Parkin quantification methods also reveal output bimodality. **(A-B)** Alternative quantifications of mitochondrial PINK1 levels or mitochondrial Parkin recruitment for single-cell data shown in Fig. 3F. n = 1987 cells. **(A)** Pathway activation estimated as maximum achieved fraction of total Parkin-mKO2 localized on mitochondria. Colors: local point density (methods). PINK1 input threshold from Fig. 3F shown as a vertical black line. **(B)** Histogram showing ability of input threshold in (A) to separate Parkin activation populations. **(C)** Pathway input estimated as PINK1 expression level. Expression levels were calculated similarly to PINK1 input scores as time-averaged values. An input threshold for this metric was recalculated and is shown as a vertical black line. **(D)** Histogram showing ability of input threshold in (C) to separate Parkin activation populations.

**Fig. S5.**
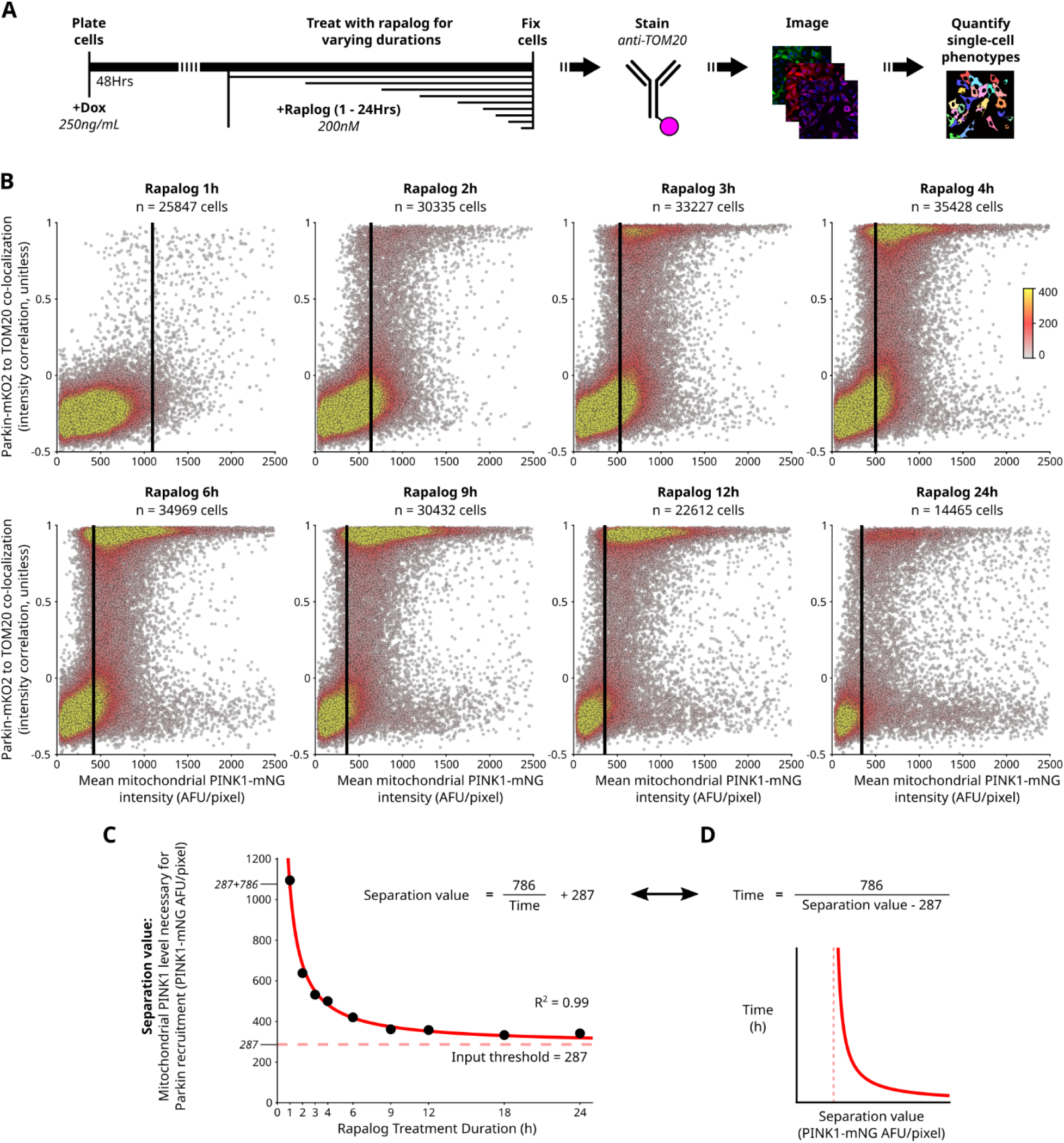
Fixed-cell data recapitulates the threshold and reciprocal delay behaviors required for activation of the PINK1/Parkin circuit. **(A)** Approach for quantifying PINK1-mNG and Parkin-mKO2 dynamics via fixed-cell timeseries. **(B)** Mitochondria-Parkin co-localization versus mitochondrial PINK1 levels for individual cells. Single-cell data from three experimental replicates was pooled. A selection of treatment durations is shown. Colors: local point density (methods). Vertical black lines denote separation values identifying the mitochondrial PINK1-mNG level capable of driving Parkin-mKO2 recruitment for that treatment duration. Relative datapoint scarcity at 24h due to mitochondrial degradation. **(C)** Calculated separation values versus rapalog treatment durations. Datapoints from (B). Fitted hyperbola, solid red line; hyperbola’s limit, dashed red line. Relationship to single cells: cells with mitochondrial PINK1 will undergo detectable Parkin translocation when the separation value drops below their mitochondrial PINK1 level. **(D)** The hyperbola, in (C), rearranged to the form used for live-cell data in Fig. 3H.

**Fig. S6.**
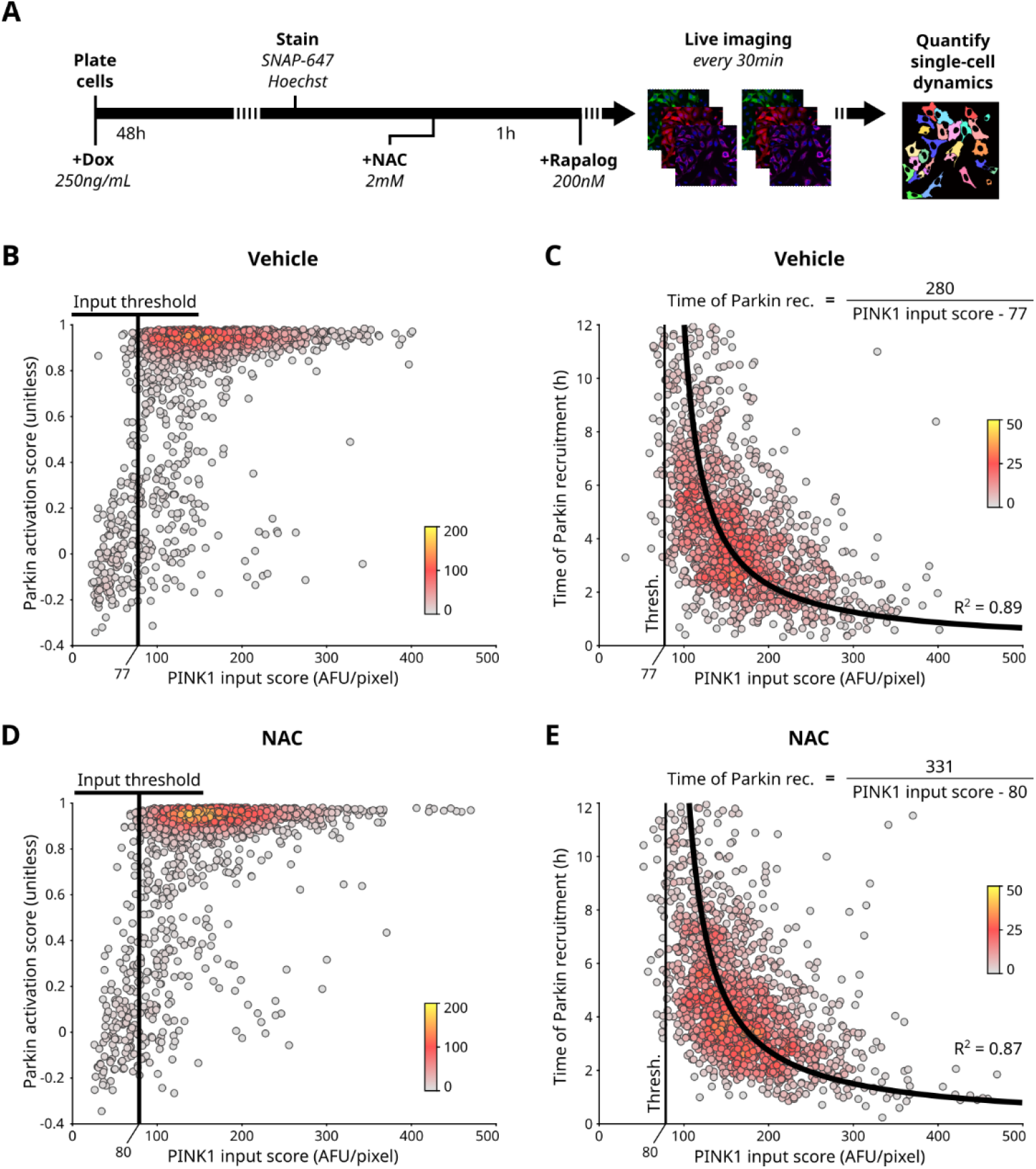
Dissecting the effects of NAC on PINK1/Parkin circuit dynamics. **(A)** Experimental schematic for measurement of PINK1/Parkin circuit dynamics in live cells following ROS modulation. All treatments were maintained through subsequent steps. **(B/D)** Parkin activation scores versus PINK1 input scores of individual cells. n = 1633 (B); n = 1721 (D). Single-cell data from two experimental replicates was pooled. Calculated threshold is marked. Colors: local point density (methods). **(C/E)** Time of Parkin recruitment vs PINK1 input scores of individual cells. n = 1375 (C); n = 1486 (E). Colors: local point density (methods). Fitted Hyperbolas are shown with R-squared values. Thresholds from (B/D) are also shown.

**Fig. S7.**
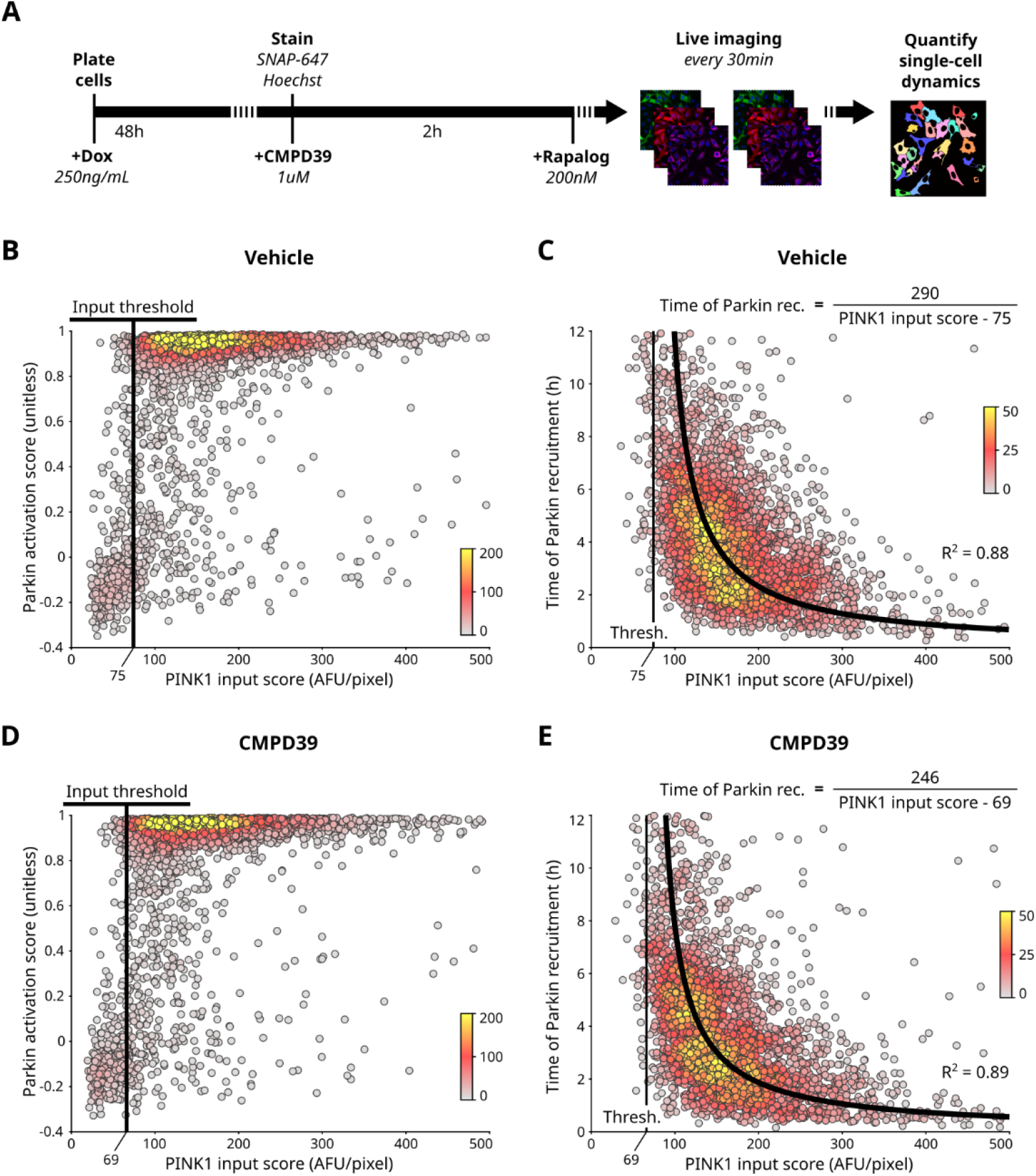
Dissecting the effects of USP30 inhibition on PINK1/Parkin circuit dynamics. **(A)** Experimental schematic for measurement of PINK1/Parkin circuit dynamics in live cells following USP30 inhibition. All treatments were maintained through subsequent steps. **(B/D)** Parkin activation scores versus PINK1 input scores of individual cells. n = 3453 (B); n = 3317 (D). Single-cell data from three experimental replicates was pooled. Calculated threshold is marked. Colors: local point density (methods). **(C/E)** Time of Parkin recruitment vs PINK1 input scores of individual cells. n = 2924 (C); n = 2794 (E). Colors: local point density (methods). Fitted Hyperbolas are shown with R-squared values. Thresholds from (B/D) are also shown.

**Figure S8:**
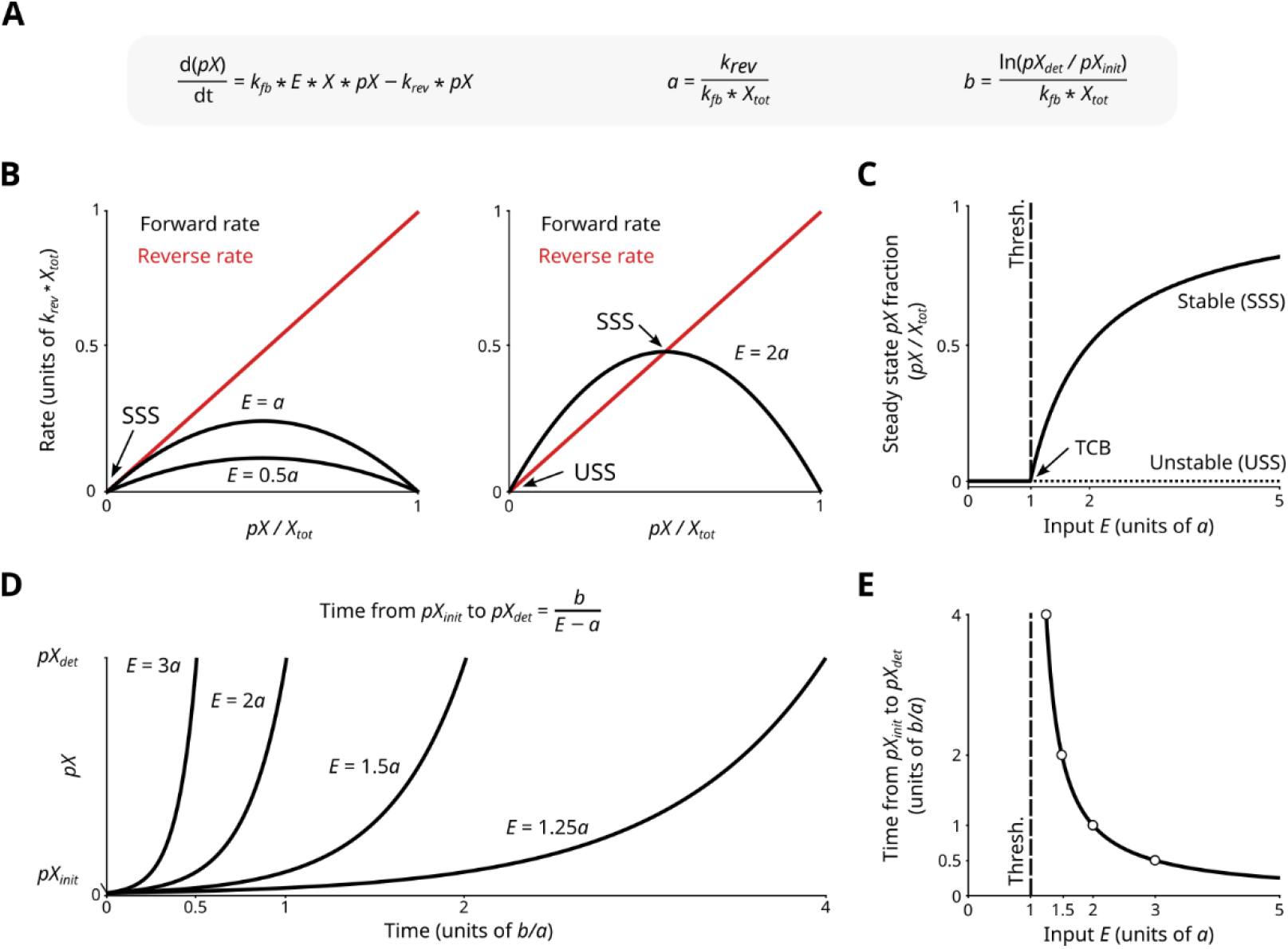
Input-coupled positive feedback produces threshold and reciprocal delay behavior (A) **(A)** Rate equations for minimal model of input-coupled positive feedback circuit as in Fig. 5A. Parameters *a* and *b* govern input threshold and reciprocal time delay, respectively. **(B)** Rate analyses for example values of *E* below or above the input threshold. Forward (*k_fb_*\**E*\**X*\**pX*) and reverse (*k_rev_*\**pX*) reaction rates are plotted. Stable steady states (SSS) and unstable steady states (USS) are identified. **(C)** Steady state analysis illustrating the system’s transcritical bifurcation (TCB) that defines the input threshold. For input *E* levels below the TCB, the system has one stable steady. For input *E* values above the TCB, the system has two stable states, one stable and one unstable. **(D)** Exponential growth of *pX* from *pX_init_* to *pX_det_* for various values of *E*. Equation describing hyperbolic relationship between *E* and time to reach *pX_det_* is shown. **(E)** Relationship between *E* and time to reach *pX_det_*. Examples from (D), marked with empty circles.

**Fig. S9.**
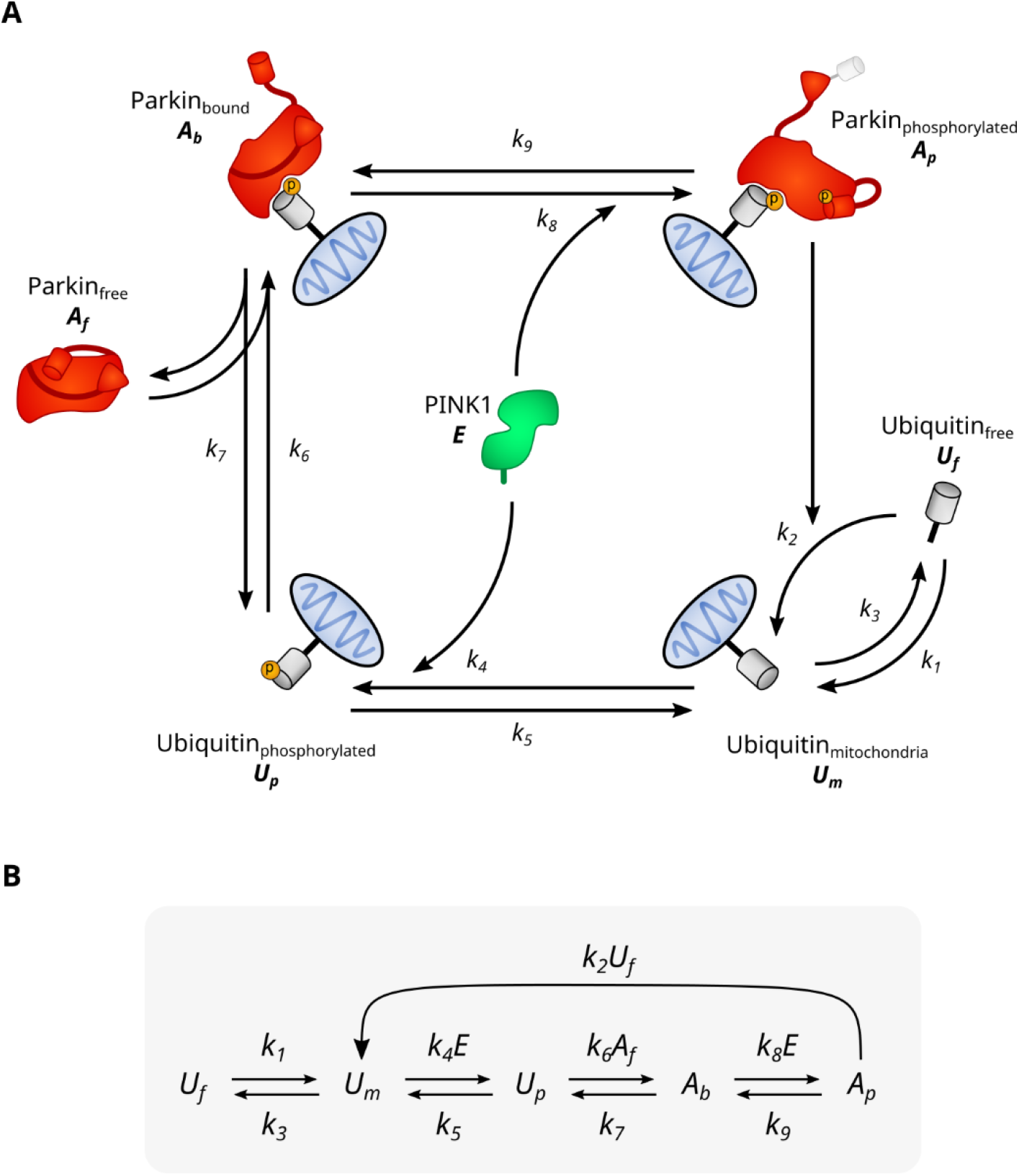
Parameterization of the PINK1/Parkin circuit for mathematical model analysis in supplemental text. **(A)** Parameterized model of the PINK1/Parkin circuit. Variables are defined: Parkin, *A;* Ubiquitin, *U;* PINK1, *E*. Numbered rate constants “*k*” are marked for each step. **(B)** Simple schematic of the model. Rate of each step obtained by multiplying the source moiety (e.g. *U_f_*) by the rate shown (e.g. *k*_1_)

## Acknowledgements

We thank all of the member of the Altschuler-Wu lab for their support. We also thank Jayanta Debnath, Orion Weiner, and Matt Jacobson for helpful feedback and perspectives during this study. We thank the UCSF HDFCCC Laboratory for Cell Analysis Shared Resource Facility for training and assistance with cell sorting on the BD FACS Aria 2. The HDFCCC Laboratory for Cell Analysis Shared Resource Facility is supported through a grant from the National Institute of Health (P30CA082103). Protein structure analyses performed via UCSF ChimeraX, developed by the Resource for Biocomputing, Visualization, and Informatics at the University of California, San Francisco. ChimeraX development supported by National Institutes of Health R01-GM129325 and the Office of Cyber Infrastructure and Computational Biology, National Institute of Allergy and Infectious Diseases. This work was funded by the National Science Foundation Graduate Research Fellowship Program 2034836 (CSW), a gift from an anonymous donor (CSW), the National Cancer Institute - National Institute of Health RO1-CA184984 (LFW) and the Chan-Zuckerberg Initiative (LFW and SJA).

## Author contributions

Conceptualization: CSW, SJA, LFW

Methodology: CSW, SBA, SJA, LFW

Investigation: CSW

Analysis: CSW

Visualization: CSW

Modeling: CSW, SBA

Funding acquisition: CSW, SJA, LFW

Supervision: SJA, LFW

Writing – original draft: CSW, SBA

Writing – review & editing: CSW, SBA, SJA, LFW

## Competing interests

SJA and LFW are founders and scientific advisory board members of Nine Square Therapeutics.

## Data and materials availability

Upon scientific review, code and processed data will be deposited on Zenodo. Due to size limitations of public repositories, raw image datasets will be made available for download by SJA and LFW upon request. Upon scientific review, sequences of all plasmids generated for this study will be deposited on Zenodo. Plasmids generated for this study will be made available by SJA and LFW upon completion of a material transfer agreement (MTA). Plasmids containing mNeonGreen additionally require an mNeonGreen user license from Allele Biotechnologies.

## Materials and Methods

### Plasmid generation

Plasmids for expressing variants of tagged Parkin: untagged Parkin (pCMV-Parkin); Untagged Parkin with E16A mutation (pCMV-Parkin(E16A)); N-terminally tagged wildtype Parkin (pCMV-mKO2-Parkin); N-terminally tagged Parkin with M1L substitution (pCMV-mKO2-Parkin(M1L)), internally tagged Parkin (pCMV-Parkin-A92mKO2), and internally tagged ligase dead Parkin (pCMV-Parkin(C431N)-A92mKO2). All above Parkin expression plasmids were derived from the pCMV-mCherry-Parkin (Addgene, #23956). The mKO2 insert was derived from pLL3.7m-Clover-Geminin(1-110)-IRES-mKO2-Cdt(30-120) (Addgene, #83841). Lentiviral Parkin expression vector (pLv-CMV-Parkin-A92mKO2), and Lentiviral tether expression vector (pLV-EF1a-SNAP-FRB-FIS1(MTS)) were created using the backbone from pLV-EF1a-IRES-Puro (Addgene, #85132). The SNAP tag insert was derived from pSNAPf (NEB, N9183S) and the FRB-Fis1 insert was derived from pC4-RhE-FRB-Fis1. The Lentiviral doxycycline-inducible FKBP-PINK1(d1-110)-mNeonGreen expression vector, pLv-TetOn-FKBP-PINK1(d1-110)-mNG-Neo, was created using the backbone from pLv-TetOn-MCS1-P2A-MCS2 (Neo) (Addgene, #89180). The FKBP insert was derived from pC4M-F2E-GFP-FKBP (Addgene, #68058), the PINK1 insert was derived from pCMVTNT-PINK1-C-Myc (Addgene, #13314) and the mNeonGreen insert was obtained as cDNA (Allele Biotech, ABP-FP-MNEONSB) (*44*). All plasmid assembly was done using standard cloning methods. All plasmids were grown following transformation into NEB STBL competent *E. coli* (NEB, C3040H). All inserts were verified via sanger sequencing (Elim Biopharmaceuticals).

### Cell culture

HeLa cells were cultured with a base media consisting of RPMI (Thermo Fisher Scientific, 11875119) with 10% fetal bovine serum (Gemini Bio., #100-106) and antibiotic/antimycotic (Thermo Fisher Scientific, 15240062). HeLa cells were chosen for this study because they lack endogenous Parkin expression (*3*, *45*). HEK293T cells were cultured with a base media consisting of DMEM (Thermo Fisher Scientific, 11965092) with 10% fetal bovine serum and penicillin/streptomycin (Corning, 30-002-CI). Immunofluorescence and live-cell imaging experiments were performed in 96 well glass bottom, tissue culture treated imaging plates (Thermo Scientific Nunc, 164588). Cells were kept at 37C and 5% CO2 during culturing and during live cell microscopy.

### Immunofluorescence

Cells were fixed in PBS (Invitrogen, AM9625) with 4% paraformaldehyde (Electron Microscopy Sciences, 15710) for 20 minutes at room temperature and were then permeabilized in PBS with 0.5% Triton X-100 (Sigma-Aldrich, 93443) for 20 minutes. Cells and then washed three times in PBS and blocked in PBS with 0.3% Triton X-100 and 3% Bovine Serum Albumin (BSA) (Fisher bioreagents, BP9703) for 1 hour. Blocked cells were Incubated in staining buffer (PBS with 0.3% Triton X-100 and 1% BSA) with primary antibody overnight at 4C when applicable. Cells were then washed three times with PBS and incubated in staining buffer with secondary antibody for 2 hours at room temperature when applicable. Fluorescently conjugated primary antibody was included in the secondary antibody stain when applicable. Finally, cells were washed twice in PBS, stained with 1:5000 hoechst 33342 (Invitrogen, H3570) for ten minutes, and washed once more in PBS. Primary antibodies: 1:500 ms-anti-Parkin (Abcam, ab77924); 1:1000 Rabbit anti-TOM20 Alexa Fluor (AF) 647 (Abcam, ab209606). Secondary antibodies: 1:1000 Goat anti-mouse AF plus 488 (Invitrogen, A32731).

### Activity assessment of Parkin tag variants

HeLa cells were seeded in 96 well imaging plates and allowed to grow for 36 hours. Individual wells were transfected with 25ng of plasmid expressing one tagged Parkin variant using a Lipofectamine 3000 kit (Invitrogen, L3000-015) via manufacturer’s instructions. Media on transfected cells was replaced after 3.5 hours and cells were allowed to recover. Wells were treated with either 20uM carbonyl cyanide 3-chlorophenylhydrazone, (CCCP; Sigma Aldrich, C2759) or 2:5000 DMSO (Sigma Life Science, D2650) control for either 10 hours or 30 minutes with timing such that the treatment ended and cells were fixed 48 hours after transfection. Cells were fixed and subjected to immunofluorescence staining for Parkin and TOM20. Cells were imaged on an Operetta automated microscope in confocal mode with a 20x water objective, no pixel binning, and standard filter sets.

### Cell line generation

Lentivirus was made using HEK293T cells and tittered for 10% transduction efficiency. HeLa cells were co-transduced with lentivirus delivering MtTether-SNAP: EF1a-SNAP-FRB-Fis1(MTS), CMV-Parkin(A92-mKO2), and TetOn-FKBP-PINK1(110-581)-mNeonGreen using 8ug/mL polybrene in growth media. Cells were selected for the TetOn PINK1 expression cassette via treatment with 400ug/mL Geniticin (Life Tech. Corp., 10131035). Triple positive cells were isolated using fluorescence-activated cell sorting (FACS). Prior to FACS, PINK1-mNG expression was induced with 50ng/mL Doxycycline Hydrochloride (Sigma-Aldrich, D3072) for 48 hours and MtTether-SNAP was stained with 1:2000 SNAP-647 SiR (NEB, s9102) for 30 minutes. Following FACS, triple positive cells were allowed to expand and recover in the absence of doxycycline, to eliminate PINK1-mNG expression prior to being frozen for storage.

### Live imaging of chemically induced recruitment of PINK1 and Parkin to mitochondria

Triple positive HeLa cells were seeded in 96 well imaging plates with 250 ng/mL doxycycline. PINK1-mNG expression was induced for 48 hours with media and doxycycline. Media with doxycycline was refreshed after 24 hours due to chemical instability of doxycycline in aqueous solution. All stain, wash, and imaging medias used in subsequent steps were growth media containing doxycycline. Prior to imaging, cells were stained with 1:1000 SNAP-AF647-SiR for 30 minutes, stained with 1:1000 hoechst 33342 for 10 minutes and washed 2×20 minutes. Hoechst staining of live HeLa cells was observed to have no effect on cell viability or growth rate. Cells were imaged in growth media once (~30 minutes) to capture the pre-treatment state. Cells were then treated with 200nM rapalog (A/C heterodimerizer, previously AP21967) (Takara Bio., 635055) and immediately subjected to live cell microscopy with images being taken every 30 minutes for 12 hours. For live cell experiments longer than 12 hours, imaging frequency was every 45 minutes. Cells were imaged using an Operetta microscope in confocal mode with a 40x water objective, 2×2 pixel binning, and standard filter sets. The exact imaging time for each was recorded.

### Modulation of threshold and rate with ROS and USP30 inhibitor

Triple positive HeLa cells were seeded, treated with Doxycycline for 48 hours, stained, and imaged as with the live cell imaging of the chemically-induced Pink1 recruitment system described above. A 1-hour pretreatment with 2mM N-acetyl cysteine (NAC; Sigma-Aldrich, A9165) was accomplished by addition to all staining, wash, and imaging medias. Vehicle treatment for NAC was water. A 2-hour pretreatment with 1uM CMPD39 (also called USP30 inhibitor 18; MedChem Express, HY-141659) was accomplished by addition to the wash and imaging medias. Vehicle treatment for CMPD39 was DMSO.

### Immunofluorescence-based analysis of induced recruitment system

Triple positive HeLa cells were seeded and treated with Doxycycline for 48 hours as with the live cell imaging of the induced recruitment system described above. Individual wells of the 96 well plates were treated with rapalog for treatment durations ranging from 1 hour to 24 hours. Treatments were started at different times such that all cells were fixed at 72 hours following seeding. Cells were fixed, subjected to immunofluorescence staining for TOM20, and imaged on an Operetta microscope in confocal mode with a 40x water objective, no pixel binning, and standard filter sets.

### Western blots

Triple positive cells were treated with 100nM Rapamycin or 200nM rapalog for either 6 or 24 hours. Cell pellets were collected and lysed in RIPA buffer (Sigma, R0278) with Halt protease and phosphatase inhibitor cocktail (Thermo Fisher Scientific, 78440). Lysates were run on Mini-Protean TGX 4-20% SDS-PAGE gels (Bio-Rad, 4561093). Western blotting was performed using standard approaches. Odyssey blocking buffer (LI-COR Biosciences, 927-50000) was used for blocking and antibody incubation steps. Western blots were imaged using a LI-COR Odyssey CLx infrared imaging system. Primary antibodies: 1:1000 Rb-anti-pS6 (Ser235/236; Cell Signaling Technologies, 4856), 1:1000 Rabbit anti-S6 (Cell Signaling Technologies, 2217), and 1:1000 Mouse anti-GAPDH (Cell Signaling Technologies, 97166). Secondary antibodies: 1:5000 Goat anti-rabbit IRDye 800CW (LI-COR, 926-32211) and 1:5000 Goat anti-mouse IRDye 680LT (LI-COR, 926-68020).

### Microscopy image processing and cell segmentation

All image quantification and analyses were done in MATLAB. Images were first subjected to flatfield illumination correction using pre-generated objective-specific flatfield correction estimations. Uniform background fluorescence was then removed for each image. Finally, mild fluorescence crossover from the TRITC channel (mKO2) into the FITC channel (mNeonGreen or AF488) was corrected in a pixelwise manner using a multiply and subtract compensation approach. Crossover correction was empirically calibrated for each microscope objective and for each set of exposure times used in this study. Correction multipliers ranged from 0.02 to 0.05. Processed images were then subjected to cell segmentation using an in-house watershed segmentation pipeline. Segmentation was able to identify nucleus and cytoplasm boundaries for each cell, in each field of view, and at each timepoint (when applicable). A perinuclear region was also identified by removing the outer 25% of pixels from the edge of the cytoplasm region in order to remove small segmentation errors due to overlap with adjacent cells.

### Single-cell quantification for Parkin tag variant comparison

Following cell segmentation, prior to quantification, single-cell data from three experimental technical replicates was pooled. For each cell, foreground images were calculated for both the anti-Parkin and anti-TOM20 channels by applying a tophat filter with a radius of 8 pixels to each cell image. Colocalization of anti-Parkin and anti-TOM20 intensity was calculated as the pixelwise Pearson correlation coefficient of the foreground images for those channels. For each cell, total anti-TOM20 intensity was calculated by identifying a mitochondrial mask (anti-TOM20 foreground image intensity above 2500 AFU) and then calculating the total pixel intensity of the original anti-TOM20 image within that mask.

To compare cells with similar Parkin expression levels, a sliding window median approach was used for each construct and each condition. Window center points were integer Parkin expression values ranging from 3000 to 20000 AFU/pixel. Window size was +/-850 AFU/pixel. Only cells with Parkin expression values between 3000 and 20000 AFU/pixel were used. For each construct, DMSO-treated data was used to normalize the CCCP-treated data. When comparing Parkin-TOM20 staining colocalization, the DMSO sliding median values were subtracted from the corresponding CCCP-treated values to estimate the change in localization across parkin expression levels. When comparing total cellular anti-TOM20 intensity, CCCP-treated values were divided by the corresponding DMSO-treated values to estimate the proportion of anti-TOM20 staining remaining across parkin expression levels. Finally, these normalized sliding medians were summarized by taking the mean value along their length. Hypothesis testing was performed using a bootstrap approach of random resampling with replacement of single-cell data, 1000 times. Due to all 6 Parkin tag/mutant variants (Untagged, Internal, Nterm-M1, Nterm-I1, LD, and Untagged-E16A) being assessed in parallel, a Bonferroni multiple comparison correction for 5 comparisons was used.

### Single-cell quantification and tracking for live-cell PINK1/Parkin dynamics

Following single-cell segmentation, foreground images and mitochondrial masks were calculated. First, foreground images were calculated for all images by applying a 2D bandpass gaussian filter (sequential high pass gaussian with a wavelength of 50 pixels to remove background intensity and low pass gaussian with wavelength of 3 pixels to reduce pixel noise) using the filt2 MATLAB function (*46*). Then, a mitochondria mask was calculated using the MtTether-SNAP foreground image and a threshold that was calculated on a per-cell basis. This cell-specific threshold was calculated as the median pixel intensity of the original cell MtTether-SNAP image raised to the exponent of 0.8 and multiplied by 2. This approach was empirically designed to be effective over a range of MtTether-SNAP expression levels.

Next, a wide variety of distinct cellular phenotypes (features) were quantified. Colocalization features were calculated as the pixelwise Pearson correlation coefficients between pairs of foreground images. Mitochondrial PINK1 levels were calculated as the mean pixelwise PINK1-mNG foreground intensity in the mitochondrial mask. Colocalization features and mitochondrial PINK1-mNG features were calculated only in the perinuclear region to avoid intensity artifacts at cell edges. Expression level features were calculated as means from the original channel intensity images.

Following feature extraction, single cells were tracked from one timepoint to the next by finding the closest cell both in distance and phenotypic similarity. This was accomplished using a distance metric defined as the weighted sum of: the distance between nucleus centroids, the ratio of cell areas, the ratio of nucleus area, and the ratio of marker expression levels between candidate cells. Only cells which were successfully tracked across all timepoints were used for downstream analyses. Additionally, cells with high or low Parkin-mKO2 expression or tether expression levels (approximately top and bottom 5%) were discarded. Cells with high PINK1-mNG expression levels above 10000 AFU/px were discarded as well due to masking of PINK1-mNG recruitment to mitochondria at such high expression levels. Similarly, cells which did not show an increased PINK1-mitochondria colocalization upon rapalog treatment were discarded as well. Quantification of mitochondrial PINK1 levels was prone to overestimation for cells with no PINK1-mitochondria colocalization, but was effective if PINK1 was colocalized with mitochondria. Finally, because some cells with low initial PINK1-mNG expression levels underwent doxycycline-dependent PINK1-mNG expression induction during the experiment, cells with PINK1-mNG expression levels which increased 1.5-fold or more over the course of the experiment were discarded.

### Quantification of PINK1 input, and detection time of Parkin recruitment, and Parkin output scores for live cell data

Each cell was quantified individually. First, the Parkin output score was calculated as the maximum observed colocalization between the Parkin-mKO2 and MtTether-SNAP channels. The maximum value was used because extended Parkin-mKO2 recruitment eventually caused mitochondrial degradation and an associated decrease in measured foreground colocalization. Next, the detection time of Parkin recruitment was determined as the time at which the colocalization of Parkin-mKO2 and MtTether-SNAP reached a value of 0.4. The value 0.4 was chosen because it is half-way between the values measured for cells with and without parkin recruitment. Finally, the PINK1 input score was calculated from the mitochondrial PINK1 level. Due to noise over time in this quantification, the data was smoothed via a moving mean with a window size of 5 timepoints. The PINK1 input score was then calculated by taking the mean value of this smoothed data from when PINK1 was recruited either to when Parkin was recruited (for cells with Parkin response scores > 0.4) or to the end of the experiment (for cells with Parkin response scores < 0.4).

### Threshold and delay function quantification from live cell PINK1/Parkin experiments

First, an input threshold value was defined by: 1) taking a sliding window over the single-cell PINK1 input scores (overall width of 5% of the population size); 2) calculating the percentage of cells within each window exhibiting parkin recruitment (Parkin output score > 0.4); and 3) identifying the center point of the first window (moving from small to large PINK1 input scores) with at least 50% of cells exhibiting Parkin recruitment. Next a hyperbolic curve was fit to the PINK1 input score vs detection time of Parkin recruitment data. This hyperbola was fit using a geometric distance, sum of squares, approach. Specifically, the above-calculated input threshold was used as a constant in the following function, and the “delay scaler” in the following function was optimized to obtain a fit: *Detection time of Parkin recruitment = (delay scaler) / (PINK1 input score – input threshold*). When comparing across conditions, hypothesis testing was performed using a bootstrap approach. Random sampling with replacement and re-quantification of the input threshold and delay scaler values was performed 1000 times. A Bonferroni multiple comparison correction was used to correct for 2 comparisons in each experiment.

### Single-cell quantification for fixed-cell PINK1/Parkin dynamics

Following cell segmentation, foreground images were calculated as for live cell data above. Gaussian bandpass wavelengths used were 100 pixels and 3 pixels. A Mitochondria mask was calculated using the anti-TOM20 mitochondrial foreground image and a fixed threshold of 1000 AFU. A single intensity threshold was used to calculate the mitochondria mask due to uniform TOM20 staining across cells. Next, features were calculated the same way as described above for live cell data.

Finally, the data quality control was performed. Mis-segmented cells (small cell areas, small nucleus areas, or high levels of hoechst staining in the cytoplasm) were discarded. Cells with high or low Parkin expression (approximately top or bottom 5%) were discarded. Cells with low TOM20 staining (average cellular intensity less than 1200 AFU/pixel) were discarded to remove cells that had undergone mitochondrial degradation at longer timepoints. Cells with a low PINK1-TOM20 intensity colocalization scores (less than 0.6) were discarded to ensure that only cells with mitochondrial PINK1 were analyzed.

### Threshold and delay function quantification from fixed cell PINK1/Parkin experiments

Each cell had a quantified mitochondrial PINK1 level (PINK1 intensity on mitochondria) and a Parkin recruitment level (Parkin-TOM20 intensity colocalization). At each timepoint (rapalog treatment length), a variant of Otsu’s thresholding method was used to identify a PINK1 level capable of separating cells into two Parkin recruitment level populations. In short, an algorithm searched for a PINK1 level separation value, able to minimize variance in Parkin recruitment level for the two populations of cells: PINK1 levels above or below the separation value. Functionally, this separation value identified the mitochondrial PINK1 level required to trigger Parkin recruitment for each length of rapalog treatment. A hyperbola with the equation *PINK1 separation value = A/(rapalog treatment time) + B* was fit to the data by identifying values of A and B using a nonlinear least squares approach. The hyperbola fitted here is an algebraic rearrangement (swapping the dependent and independent variables) of the hyperbola fit to the live cell data. The hyperbolic relationship extracted from the fixed cell data (required PINK1 level as a function of Rapalog treatment time) is distinct in identity but similar in nature to the hyperbolic relationship extracted from the live cell data (Parkin recruitment time as a function of Mitochondrial PINK1 levels).

### Coloring datapoints by local point density

Local point density was calculated by counting the number of nearby points within a radius of 5% of the field of view in each direction (e.g., if the x-axis range is [0,200], then the averaging field around a point x is x ± 10).

### Simulation of heterogeneity for minimal model of input-coupled positive feedback

To simulate heterogeneity, we simulated 2000 “cells”. For each cell, values for *E*, *k_rev_*, and *pX_init_* were sampled as follows.

*E* values were randomly sampled using a gamma distribution (shape parameter: 5.14; scale parameter: 0.38). The gamma distribution was empirically fit to the observed distribution of Pink1 input scores using MATLAB’s “fitdist” function.

Values for *k_rev_* were sampled from a log normal distribution (mu: 0; sigma: 0.5). Values of *pX_init_* were scaled by dividing with the same random *k_rev_* multiplier to reflect the expected relationship between the two parameters (e.g. deubiquitinases govern baseline levels of mitochondrial ubiquitin (*6*, *16*, *17*, *35*)).

The detection level was set to be *pX_det_=100*pX_init_*, and the steady state *[pX]/[X_tot_]* and time to detectable activation were solved analytically.

### Plotting algebraic solutions for minimal model of input-coupled positive feedback

Algebraic solutions for steady state *pX/X_tot_* and time to *pX_det_* (solid lines in Fig. 5B-C and Fig. S8C,E) were plotted using the following equations: 1) *Steady state*=1-1/*E* with negative steady state values set to zero, and 2) *time to pX_det_*=1/(1-*E*). In Fig. S8B, reverse rates were plotted as *rate*=*pX/X_tot_*, and forward rates were plotted as *rate*=*pX/X_tot_*-(*pX/X_tot_*)^2^. In Fig. S8D, exponential growth curves were plotted as *pX*=*p_Xinit_ e*^((*E*-1)\**time*\**ln*(*pX_det_/pX_init_*)). Values of *pX_det_*=0.1 and *pX_init_*=0.001 were used for plotting, though any pair of values *pX_det_* < *pX_init_* would also work, with the only change being where where *pX_init_* lies along the y-axis.

### Analysis of existing UBL domain structures

PDB accession numbers for structures shown in Fig. S2 are as follows: hParkin, 5c1z; hUbiquitin, 1f9j; hRAD23B, 1uel; hNedd8, 1ndd. Visualization and analysis were performed in the UCSF ChimeraX software (*47*). Hydrogen bonds and salt bridges were predicted using the show hydrogen bonds option in UCSF ChimeraX.

### Conservation at the N-terminus of N-terminal ubiquitin-like domains

The amino acid sequence and the metadata for 67594 annotated UBL domains in the Uniprot database were downloaded (April 4, 2019). Analysis performed in MATLAB. Entries with annotation errors were identified and discarded as follows. UBL domain entries with non-standard residues (including “J”, “O”, “U”, “X” or “Z”) or with lengths of less than 65 amino acids were discarded. Domains annotated to start at the protein’s N-terminus, but which did not start with methionine were discarded. Entries corresponding to Ubiquitin polymers rather than to UBL domains were identified as starting or ending with tandem glycines. These Ubiquitin entries were discarded to prevent over-representation in the dataset. Finally, any additional UBL domains originating from proteins containing at least one discarded domain were also discarded. This cleanup left 28447 UBL domain entries.

Alignment using traditional methods could not be used successfully due to the large number of sequences and high observed sequence variability. Instead, the conserved positioning of specific residues in the stereotyped ubiquitin fold were used to align the sequences. Specifically, the following schema was used: hydrophobic residues in the first beta sheet at positions 1, 3 and, 5; a variable-length linker with length “*x*” ranging from 7 to 17 residues; hydrophobic residues in the second beta sheet at positions *5+x+1* and *5+x+3;* hydrophobic residues in the first alpha helix at positions *5+x+9, 5+x+12*, and *5+x+16*. These positions correspond to M1, V3, V5, V15, V17, I23, L26 and V30 of Parkin’s UBL domain. Potential alignments were evaluated using scoring system where non-standard hydrophobic residues were allowed but were assigned a penalty score. Furthermore, extreme linker lengths, “*x*”, were penalized for being too short or too long. Finally, a small tie-breaker bonus was awarded to alignments which contained a positively charged residue at the equivalent position of K27 of Parkin’s UBL. This positively charged residue was used in the alignment because it was present in nearly all of the published UBL domain structures which were used as reference when designing the alignment schema. Using this approach, 19623 sequences out of 28447 were successfully aligned for residues surrounding the region of interest, M1 and E16 of Parkin’s UBL.

Finally, aligned sequences were separated into those belonging to N-terminal or internal UBL domains and were submitted to WebLogo3 (*48*) to create the sequence logos used to visualize residue conservation.

### Alignment of mammalian Parkin ortholog protein sequences

Full protein sequences for Parkin were downloaded from Uniprot for human (O60260) pig (Q2L7G3), dog (A0A8C0PPD2), rat (Q9JK66), mouse (Q9WVS6), and guinea pig (H0V739). These sequences were aligned using the Clustal Omega multiple sequence alignment tool by EMBL-EBI (*49*).

## Supplementary text

### MATHEMATICAL MODEL ANALYSIS

#### General framework

##### Assumptions and conclusions

We consider two models of increasing complexity for the dynamics in the system describing Pink1–Parkin–Ubiquitin in a Mitochondrion discussed in the main body of the paper. Simplifying somewhat, we may say that in the experiment the amount *P*(*t*) at time *t* of Parkin in a mitochondrion is tracked as it grows until it reaches a certain threshold level *P*_th_. The initial amount of Parkin *P*_in_ is assumed to be much smaller than the threshold level *P*_th_. This yields a measurement of the time *T* it takes the Parkin level to grow from *P*_in_ to *P*_th_. The level *E* of Pink1 affects the rate at which Parkin increases and thus the experiment gives us a measurement of the growth time T as a function of the Pink1 level E.

The main assumptions concerning the system that we make are

i. The system contains a positive feedback mechanism
ii. Pink1 acts as a catalyst: some of the reaction rates in the system increase when the amount of Pink1 is increased
iii. The initial amounts of Parkin and Ubiquitin on the Mitochondria are much smaller than the threshold amount
iv. The rate at which free Ubiquitin spontaneously moves to the mitochondrion so small that it may be ignored
v. The amounts of free Ubiquitin and Parkin are large and may be considered constant

These assumptions lead to the following consequences:

###### Exponential Growth

The amount of Parkin grows (or decays) exponentially; if *P*(*t*) is the amount of Parkin at time *t*, then the models predict *P*(*t*) ≈ *P*_in_*e*^*λ_E_t*^, where the growth rate *λ_E_* depends on the many reaction rates in the model, and in particular on the amount E of Pink1 in the system.

###### Switch-like behavior

The system exhibits switch-like behavior in its dependence on E: there is a critical value *E*_*_ such that *λ_E_* < 0 when *E* < *E*_*_ and such that *λ_E_* > 0 when *E* > *E*_*_. This means that if the Pink1 amount is below *E*_*_ the Parkin levels will decrease exponentially, and if the Pink1 amount exceeds *E*_*_ then Parkin grows exponentially.

###### Time to detection decreases with increasing Pink1 levels

The exponent *λ_E_* is an increasing function of the Pink1 level *E*. When *λ_E_* > 0 the Parkin level grows exponentially according to

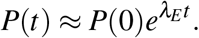

Let *P*_th_ be the threshold level at which Parkin is detected, and assume that the initial amount *P*_in_ of Parkin is small compared to *P*_th_. If the growth rate *λ_E_* is negative then the amount *P*(*t*) of Parkin will only decay and thus never reach the threshold level. On the other hand, if the growth rate *λ_E_* is positive, then the threshold time *T*_th_(*E*) follows from

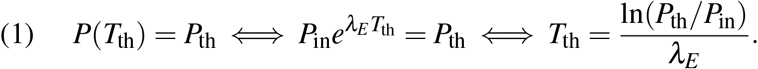

For each of the models we find that the growth rate *λ_E_* increases when *E* is increased, which therefore implies that the time to detection *T*_th_ is a decreasing function of *E*.

##### General form of the models

In each of the models we have a vector

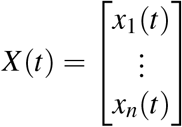

containing the total amounts of the components that the model tracks, such as Parkin, Ubiquitin, (phosphorelated or not), or combinations of these. Using mass action kinetics we arrive at a system of differential equations governing the time dependence of *X*(*t*). Assuming that we only consider the system when the amounts of non-free Parkin and Ubiquitin are small, we arrive at a linear system

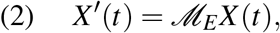

where

- E is the amount of Pink1 in the system; this quantity is assumed to be kept constant;
- 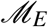 is a matrix containing the reaction rates, and which can be further decomposed as

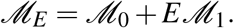 The matrix 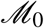 contains the rates at which reactions take place in the absence of Pink1, while 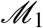 accounts for the change in the reaction rates caused by the presence of Pink1.

##### Eigenvalue analysis of the models

Using the method of eigenvalues and vectors one shows that the general solution of a linear equation such as (Eq. 2) is a superposition of exponentially growing or decaying terms, i.e.

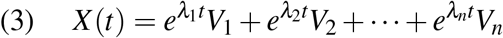

in which *λ*_1_,…,*λ_n_* are the eigenvalues of the matrix 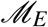, *V*_1_,…,*V_n_* are corresponding eigen-vectors. Since we are interested in the time it takes *X*(*t*) to grow from a small initial amount to a detectable threshold value, we want to consider the fastest growing term(s) in (Eq. 3), i.e. the terms corresponding to the largest eigenvalues *λ_i_*. In studying the eigenvalues *λ*_1_,…, *λ_n_* we note that in all versions of our model the matrix 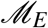 has non-negative off-diagonal entries, and is <u>irreducible</u>. This implies, by the Perron-Frobenius theorem, that 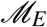 has a unique dominant eigenvalue *λ_E_*, i.e.

i. *λ_E_* is a real eigenvalue of 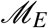 (not a complex eigenvalue)
ii. every other (possibly complex) eigenvalue *μ* of 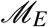 satisfies Re*μ* < *λ_E_*
iii. corresponding to the eigenvalue *λ_E_*, the matrix 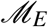 has positive left and right eigenvectors *W_E_, V_E_* respectively; by definition these satisfy

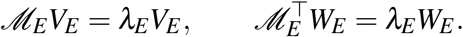 They can be normalized so that 〈*W_E_, V_E_*〉 = 1.

The dominant eigenvalue tells us the largest exponential rate with which solutions to (Eq. 2) can grow. More precisely, the eigenvalue decomposition (Eq. 3) contains one term corresponding to the dominant eigenvalue *λ_E_*. If we denote this term by *x*(*t*)*V_E_* and group the remaining terms into a slower growing component *Y*(*t*) then we have

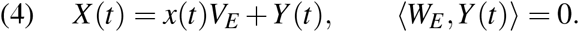

The left-eigenvector *W_E_* allows one to find the coefficient *x*(*t*) from the vector *X*(*t*) via

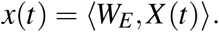

By taking the inner product with *W_E_* on both sides in (Eq. 2) one finds that the *V_E_* component of *X* satisfies an ordinary differential equation

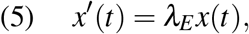

whose solution can be written as

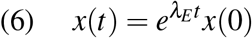

To compute the time to detection we assume that Parkin is detected when *x*(*t*) reaches a specific threshold value *x*_th_. Then the time to detection is

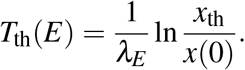

#### The minimal Pink-Parkin model

In our simplest model we only keep track of the Parkin in the system, assuming it exists in one of two states: on the mitochondria or free (not on the mitochondria).

Parkin on the mitochondria can bind free Parkin and this process is aided by Pink1 as an enzymatic catalyst. Parkin on the mitochondria also spontaneously leaves the mitochondria.

Our model keeps track of the following quantities:

*A_t_* total amount of Parkin in the system; a constant
*A_m_* amount of Parkin on the mitochondria
*A_f_* amount of free Parkin, *A_f_* + *A_m_* = *A_t_*
*E* amount of Pink1 in the system; constant in time

Since *A_t_* and *E* are time independent and since *A_m_* and *A_f_* are constrained by *A_m_* + *A_f_* = *A_t_*, the time evolution of the system is completely determined by that of *A_m_*. The following differential equation takes both recruitment and degradation into account:

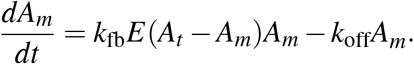

Here *k*_fb_ and *k*_off_ are reaction constants.

We make one further simplifying assumption, namely, during the observations in the experiment, the total amount *A_t_* of Parkin is much larger than the amount *A_m_* on the mitochondria. We may therefore replace *A_t_* – *A_m_* by *A_t_*, which leads us to the differential equation

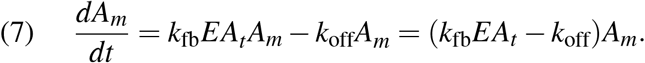

The coefficient *k*_fb_*EA_t_* – *k*_off_ is constant in time, so this differential equation is of the type 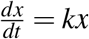, and its solution is given by the exponential growth formula *x*(*t*) = *e^kt^x*(0). In terms of *A_m_* we get

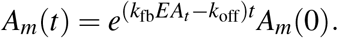

Indeed, equation (Eq. 7) is of the form (Eq. 2), if lets *X*(*t*) be the vector with only one component *X*(*t*) = [*A_m_*(*t*)], and if one lets 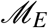 be the 1 × 1-matrix 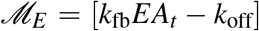. The dominant (and only) eigenvalue of 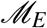 is *λ_E_* = *k*_fb_*EA_t_* – *k_off_*. However, since both the vector *X*(*t*) and the matrix 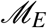 only have one component the eigenvalue analysis is not needed to solve the differential equation (Eq. 7).

In the experiment one begins with a given small amount *A_m_*(0) of Parkin and measures how long it takes before the amount *A_m_*(*t*) of Parkin reaches a fixed detectable level, *A*_th_. By solving the equation *A_m_*(*t*) = *A*_th_ for *t* we find

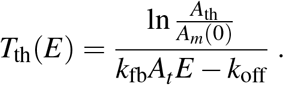

The critical value of E at which the systems “switches on” is

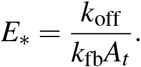

#### The full model

##### The nonlinear model

The following variables appear in the full model

- *U_f_, U_m_, U_p_* The free, mitochondrial, and phosphorelated forms of Ubiquitin
- *A_f_*, *A_b_*, *A_p_* The free, bound, and phosphorelated and bound forms of Parkin
- *E* The amount of Pink1 present in the system; a constant.

Assuming mass-action kinetics, these variables evolve according to the following set of differential equations (see model schematic in Fig. S8B).

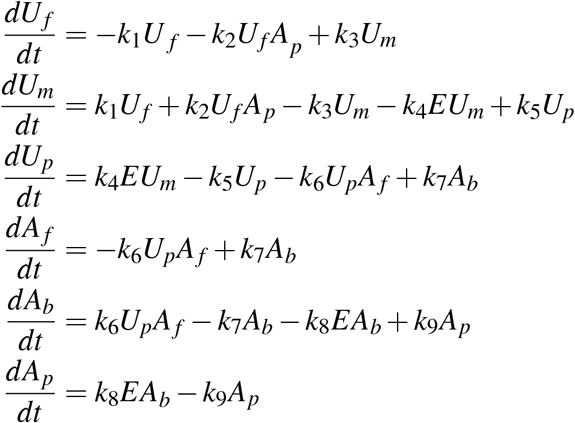

##### Linearization assuming abundant free Parkin and Ubiquitin

We can simplify the model by assuming that *A_f_* and *U_f_* are nearly constant because free Parkin and Ubiquitin are abundantly present. This leads to a reduced system with four components *U_m_, U_p_, A_b_,A_p_*, which satisfy the following four linear differential equations:

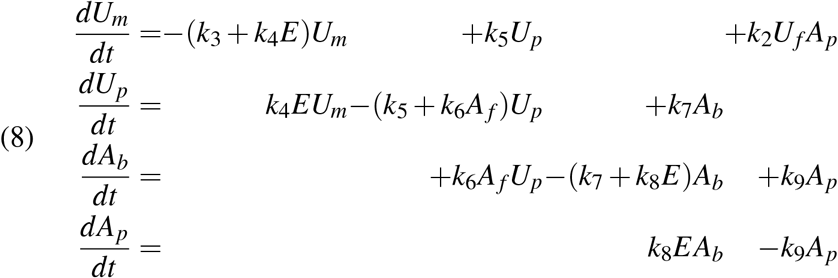

This linear system is of the form 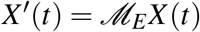 (see (Eq. 2)) where

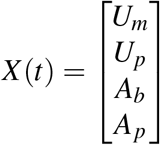

The matrix in this linear system is

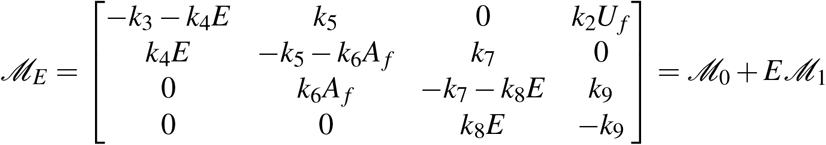

where

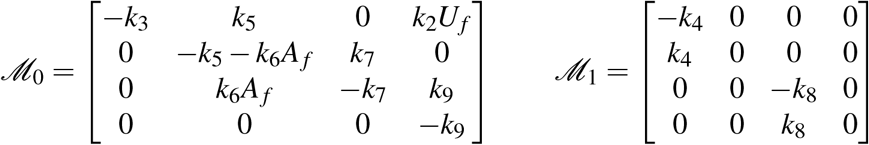

The dominant eigenvalue of 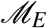 is the largest real root of the characteristic equation

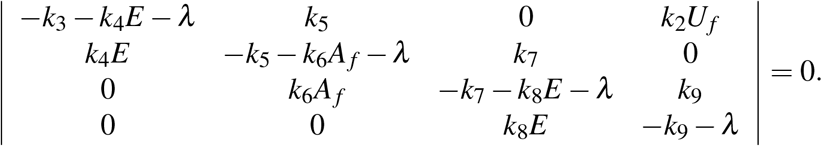

###### Theorem.

Assume *k*_2_, *k*_4_, *k*_5_, *k*_6_, *k*_7_, *k*_8_, *k*_9_, *U_f_*, and *A_f_* are positive constants. Then

a. The dominant eigenvalue *λ_E_* is a continuous function of *E* for *E* ≥ 0.
b. When *E* = 0, one has *λ_E_* < 0
c. For *E* → ∞ the dominant eigenvalue converges to a positive limiting value 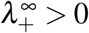.
d. There is a unique positive number *E*_*_ such that *λ*_E_*__ = 0 and for all *E* ≥ *E*_*_ one has 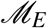.

<u>Proof.</u> The dominant eigenvalue is a simple eigenvalue and therefore depends continuously on the parameters in the matrix 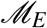. Statement (a) follows.

To verify (b), we set *E* =0 and compute

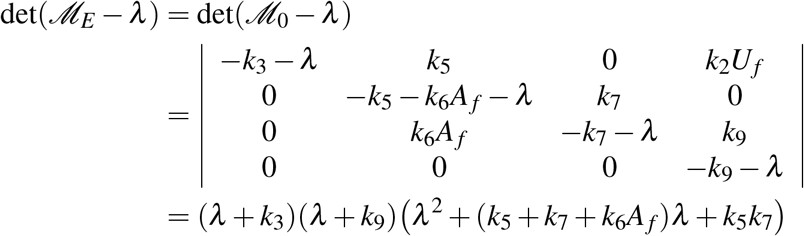

It follows that when *E* = 0 the eigenvalues of the matrix 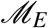 are

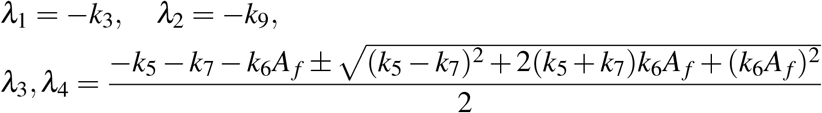

All four eigenvalues are real and negative. The dominant eigenvalue *λ_E_* is the largest of these,

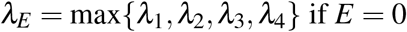

and it follows that for *E* = 0 one has *λ_E_* < 0.

We turn to (c). To analyze the eigenvalues for large values of *E* we write the characteristic polynomial as

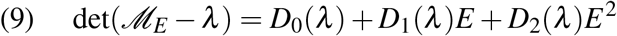

where *D*_0_, *D*_1_, *D*_2_ are polynomials in λ, which upon computation turn out to satisfy

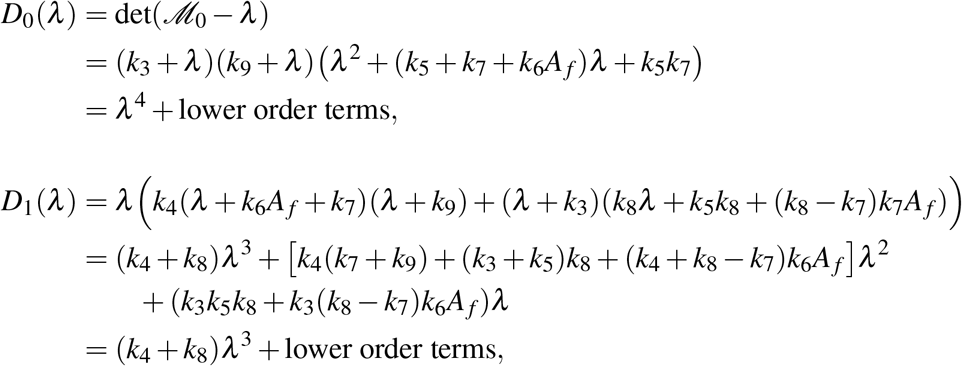

and

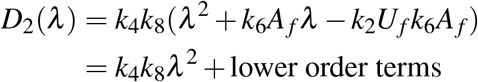

For large values of *E* the four eigenvalues of 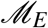 can be separated into those eigenvalues *λ* for which *λ* ≪ *E*, and those for which *λ* is comparable to *E* or larger.

If |*λ*| ~ *E* or |*λ*| » *E* then the dominant terms in the characteristic polynomial (Eq. 9) are those that contain *λ*^4^, *λ*^3^*E*, *λ*^2^*E*^2^. Two eigenvalues are therefore approximated by the nonzero roots of

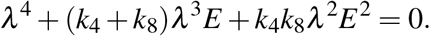

This yields two very negative eigenvalues

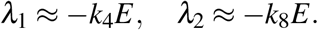

If on the other hand |*λ*| ≪ *E* then *D*_2_(*λ*)*E*^2^ is the dominant term in the characteristic polynomial (Eq. 9), and thus two of the eigenvalues are close to the roots of *D*_2_(*λ*) = 0, i.e. *λ*^2^ + *k*_6_*A_f_λ* – *k*_2_*U_f_k*_6_*A_f_* = 0, which are given by

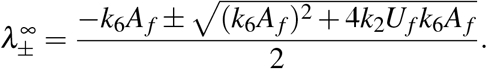

Of these, 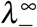 is negative and 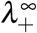 is positive. Since 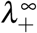 is the only positive eigenvalue, it is the dominant eigenvalue.

The dominant eigenvalue therefore satisfies

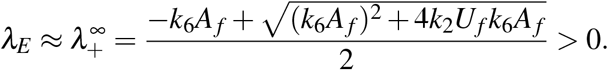

This implies statement (c).

We conclude by verifying (d). Since *λ_E_* is a simple eigenvalue, it is a differentiable function of *E* and its derivative is given by

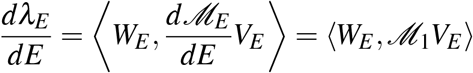

where *W_E_* and *V_E_* are the left and right eigenvalues normalized by 〈*W_E_, V_E_*〉 = 1.

If we write

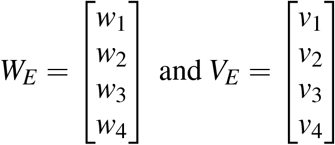

then

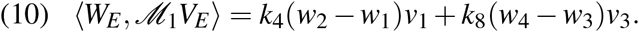

Since *v*_1_, *v*_3_ > 0 it suffices to show that *w*_2_ > *w*_1_ and *w*_4_ > *w*_3_. Expanding the eigenvalue equation 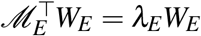 and rearranging terms we get

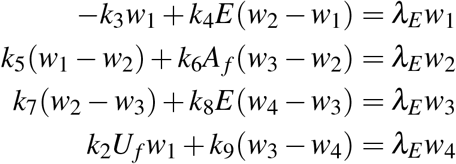

Using the assumption that *λ_E_* ≥ 0 we conclude from the first equation

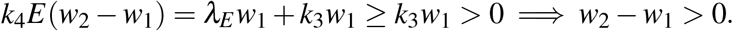

The second equation then implies

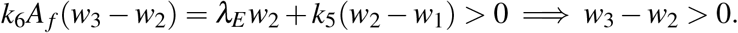

Finally the third equation leads to

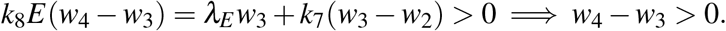

It follows that the terms on the right in (Eq. 10) are positive, and hence 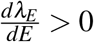.

To complete the proof we note that we have shown that the continuous function *E* ↦ *λ_E_* satisfies *λ*_0_ < 0 and *λ_E_* > 0 for all large enough *E*. Therefore there is an *E*_*_ > 0 at which *λ*_*E*\*_ = 0. Since *λ_E_* is strictly increasing when *E* > 0 there cannot be two different values *E*_*_, *E*_**_ at which *λ*_*E*\*_ = *λ*_*E*\*\*_ = 0.

## Notes

### Competing Interest Statement

Steven J. Altschuler and Lani F. Wu are founders and scientific advisory board members of Nine Square Therapeutics.

### Summary of Updates

Funding information in acknowledgements section updated

